# Trends in Machine Learning and Feature Selection Stability for Human Gut Microbiome (Shotgun Metagenomics) and Metabolomics Matched Datasets

**DOI:** 10.1101/2025.06.21.660858

**Authors:** Suzette N. Palmer, Animesh A. Mishra, Christina M. Zarek, Shuheng Gan, Ruheng Wang, Jiwoong Kim, Dajiang Liu, Andrew Y. Koh, Xiaowei Zhan

## Abstract

Microbiome research is often limited by methodological inconsistencies that reduce reproducibility and functional insight. Traditional taxonomy-based profiling is limited by sparse data, variable resolution, and reliance on gDNA sequencing which provides only indirect links to microbial function. Multi-omics integration offers a framework for linking community composition to functional outputs, but progress has been hindered by the lack of standardized frameworks and inconsistent use of machine learning. In particular, feature selection stability, which is central to biomarker discovery and experimental validation, remains underexplored. Here, we systematically benchmarked three widely used algorithms (Elastic Net, Random Forest, XGBoost) across seven multi-omics integration strategies and single-omics models. Additionally, we evaluated the impact of transforming metabolomics and taxonomic abundance data. Using human gut microbiome datasets that integrate metagenomic taxonomic profiles with metabolomics, we evaluated models for 9 binary and 8 continuous outcomes across 20 train/test splits per dataset. We further assessed the effect of feature reduction on both predictive accuracy and feature selection stability. Nonlinear learners were most consistently competitive: continuous outcomes favored metabolomics-dominant models, whereas binary outcomes favored stacked multi-omics models. Random Forest and XGBoost also yielded greater feature selection stability, particularly for full dimensional metabolomics data. Together, these findings demonstrate how integration strategy, algorithm choice, and data preprocessing jointly shape predictive performance and feature selection reproducibility in multi-omics microbiome modeling.

## Introduction

The gut microbiome plays a critical role in host homeostasis and has been implicated in multiple diseases [1,2]. However, much human microbiome research remains observational, and reproducibility across studies is limited by biological heterogeneity, analytical variation, methodological inconsistencies and data-specific challenges [3–5]. Most microbiome studies focus on taxonomy-based profiling (e.g., 16S rRNA sequencing, metagenomic shotgun sequencing), which introduces challenges such as 1) data sparsity (i.e., a large proportion of zeros representing absent taxa); 2) varying levels of taxonomic resolution; and 3) reliance on DNA-based profiling, which does not necessarily reflect functional microbial activity, such as metabolite or protein production [6,7]. Together, these limitations hinder efforts to move beyond association-based findings to mechanistic insights.

Multi-omics integration has become a promising strategy for linking taxonomic profiles with functional omics data to uncover disease-relevant microbial interactions [8–10]. However, there is substantial variation across microbiome studies in data processing, resolution, and integration approaches [11–16]. The absence of standardized frameworks may contribute to inconsistent findings in microbiome research [17–19]. Differences in machine learning models and multi-omics integration strategies can yield markedly different results, complicating the identification of reproducible taxa–function associations and the validation of disease-related patterns across studies [10,20,21]. Evaluating these models therefore requires consideration of two complementary properties: relative predictive performance and feature selection stability. Relative predictive performance describes how well an integration strategy predicts a clinical outcome compared with alternative strategies evaluated under the same validation framework. Feature selection stability describes the extent to which repeated analyses identify the same taxonomic and functional features. Together, these properties show whether an integration strategy predicts accurately and identifies reproducible patterns across model fits and data partitions. Although recent work has systematically benchmarked methods for global association testing, data summarization, individual association testing, and inter-omics feature selection in microbiome and metabolomic data [22], the relative predictive performance and feature selection stability of supervised integration strategies across clinical outcomes remains unclear.

To address these gaps, we systematically evaluated predictive performance and feature selection stability across multiple multi-omics integration approaches. Specifically, we analyzed four human gut microbiome multi-omics datasets with paired metagenomic and metabolomics data and evaluated predictive performance for binary and continuous outcomes using 20 train/test splits per dataset. We applied Random Forest, XGBoost, and Elastic Net models to full dimensional and feature reduced datasets. To assess the effect of dimensionality reduction, we filtered features within each omics dataset and compared model performance between feature-reduced and full dimensional datasets [11,14,23,24]. To evaluate the impact of scaling on predictive performance, we applied log scaling to metabolomics and centered log-ratio (CLR) transformation to taxonomic abundance data and compared these results with those from unscaled feature reduced and full dimensional datasets. We then compared model performance across single-omics and multi-omics approaches, focusing on two integration frameworks: concatenation and stacked generalization. For concatenation, we evaluated three variants: unscaled concatenation, z-score scaled concatenation, and z-score scaled concatenation with block weighting. For stacked generalization, we evaluated simple averaging, weighted Non-Negative Least Squares (NNLS) [25], elastic net (ENET) [26], and partial least squares (PLS) [27]. We then compared the performance of concatenation, stacked generalization, and single-omics models. Finally, we examined feature selection stability across machine learning models and integration strategies, comparing single-omics and concatenated approaches.

## Results

### Creation of an integrative analysis pipeline to analyze continuous and binary datasets

We collected and processed four human clinical multi-omics datasets: the Yachida colorectal cancer cohort, the Franzosa inflammatory bowel disease cohort, the Erawijantari gastrectomy cohort, and the Wang renal failure cohort. Each dataset included matched taxonomic abundance profiles generated by shotgun metagenomic sequencing and metabolomic profiles. We evaluated nine binary outcomes and eight continuous outcome analyses involving six clinical variables, selected according to the clinical characteristics of each cohort (**Table 1**) [11,14,23,24]. The binary outcomes compared the primary clinical group in each cohort (colorectal cancer, inflammatory bowel disease (IBD), post-gastrectomy status, or renal failure) with the corresponding control group. For the Yachida colorectal cancer dataset, we additionally evaluated stage-specific outcomes by comparing each cancer stage with controls. For the Franzosa dataset, we developed separate models comparing controls with the full IBD cohort and with the ulcerative colitis (UC) and Crohn’s disease (CD) subgroups. The continuous outcomes were selected from clinically relevant covariates within each cohort. For the gastrectomy cohort (Erawijantari), glucose and total cholesterol were included as markers of postoperative metabolic status, as gastrectomy is known to alter glucose homeostasis and broader lipid metabolism through changes in gastrointestinal anatomy, nutrient handling, and host metabolic regulation [28–30]. These metabolic changes have also been associated with shifts in gut microbiome and metabolome profiles following gastrectomy [23,31]. For the Franzosa cohort, fecal calprotectin (Fp) was evaluated as a continuous outcome within the control versus IBD, control versus UC, and control versus CD analysis subsets because it is a well-established marker of intestinal inflammation and is widely used to assess IBD activity [11]. For the Wang renal failure cohort, creatinine, estimated glomerular filtration rate (eGFR), and urea were included as clinical indicators of kidney function and disease severity. Renal dysfunction has also been associated with characteristic alterations in both the gut microbiome and metabolome [32,33].

**Table 1:** Summary of Input Datasets for Multi-Omics Gut Microbiome Integration Framework.

| Dataset | Outcome | Sample No. | Metabolite No. | Taxa No. |
| --- | --- | --- | --- | --- |
| Erawijantari (Gastrectomy) | Control vs. Gastrectomy | 96 (54/42) | 524 | 3103 |
|  | Total Cholesterol | 82 |  |  |
|  | Glucose | 84 |  |  |
| Franzosa (CD) | Control vs. CD | 144 (56/88) | 8848 | 5137 |
|  | Fecal Calprotectin | 107 |  |  |
| Franzosa (IBD) | Control vs. IBD | 220 (56/164) |  |  |
|  | Fecal Calprotectin | 153 |  |  |
| Franzosa (UC) | Control vs. UC | 132 (56/76) |  |  |
|  | Fecal Calprotectin | 88 |  |  |
| Wang (Kidney Failure) | Control vs. Kidney Failure | 287 (67/220) | 276 | 5293 |
|  | Creatinine | 281 |  |  |
|  | eGFR | 281 |  |  |
|  | Urea | 281 |  |  |
| Yachida (CRC) | Control vs. CRC & MP | 317 (127/190) | 450 | 5770 |
|  | Control vs. Early Lesions (MP & Stage 0 CRC) | 194 (127/67) |  |  |
|  | Control vs. Early/Mid CRC (Stages I–II) | 196 (127/69) |  |  |
|  | Control vs. Advanced CRC (Stages III–IV) | 181 (127/54) |  |  |

IBD = Inflammatory Bowel Disease; CD = Crohn's Disease; UC = Ulcerative Colitis CRC = Colorectal Cancer; MP = Multiple Polypoid Adenoma

We then developed an integrative analysis pipeline to evaluate the predictive performance of three machine learning algorithms (Elastic Net, Random Forest, XGBoost) across single-omics and multi-omics settings. Predictive performance was assessed for both binary and continuous outcomes using 20 train/test splits per dataset (**Fig. 1A-B**, **Table 1**) [11,14,23,24]. To examine the effect of dimensionality reduction, we compared models trained on feature reduced and full dimensional datasets. We also evaluated the impact of feature scaling by applying log transformation to metabolomics data and centered log-ratio (CLR) transformation to taxonomic abundance data and compared these results with those obtained using unscaled feature reduced and full dimensional datasets, respectively (**Fig. 1A**).

**Figure 1.**
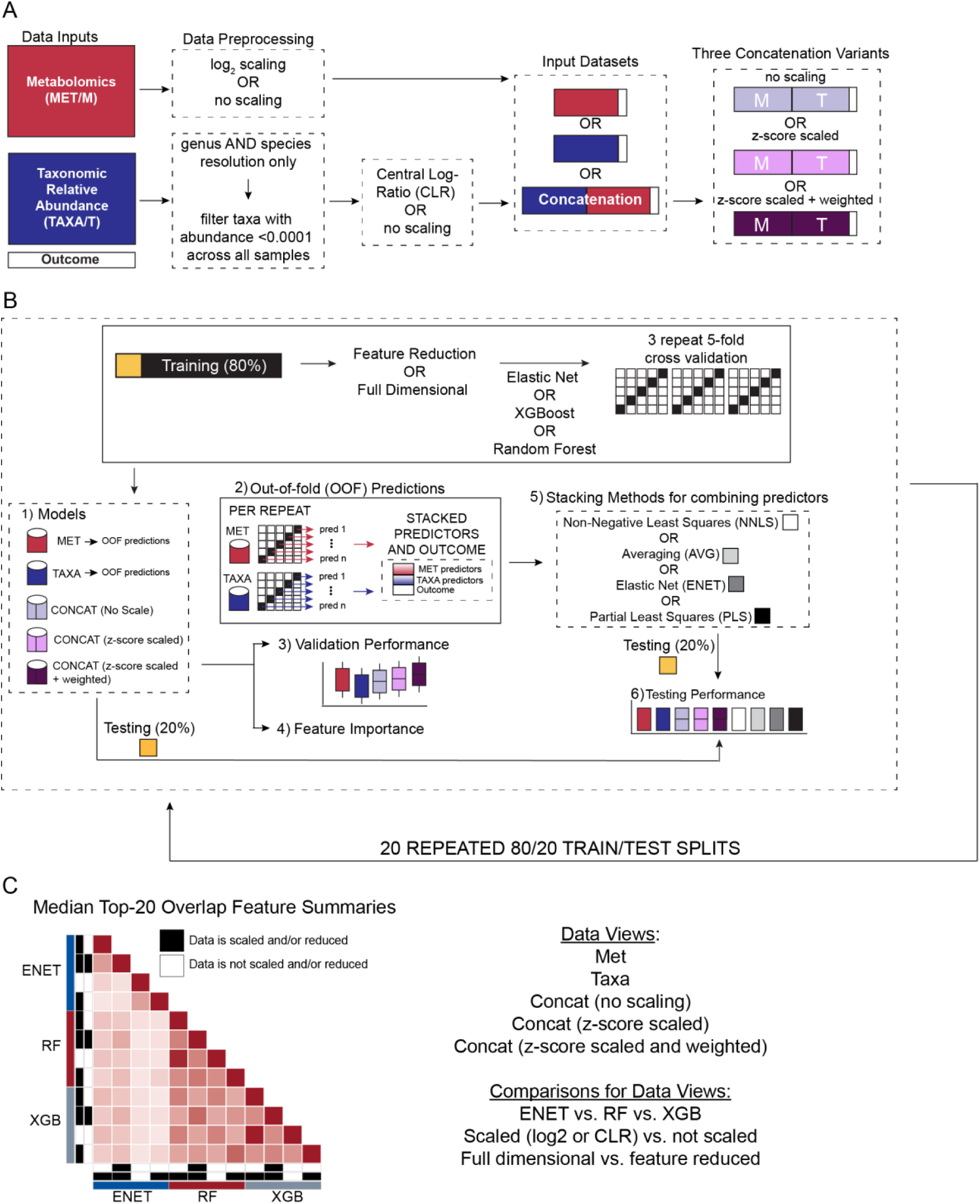
Overview of the integrative multi-omics machine-learning and feature stability pipeline. **(A)** Matched metabolomics (MET/M), taxonomic relative abundance (TAXA/T), and outcome data were used to generate single-omics and concatenated models. Metabolomics features were analyzed with or without log_2_ transformation. Genus and species-level taxonomic profiles were combined. Taxa with an abundance below 0.0001 across all samples were filtered, and the remaining features were analyzed with or without centered log-ratio (CLR) transformation. The resulting input datasets comprised metabolomics-only, taxonomy-only, and concatenated (metabolomics plus taxonomy data). For concatenated analyses, the omics blocks were combined and then evaluated without additional post-concatenation scaling, after z-score scaling, or after z-score scaling with block weighting. **(B)** Each dataset was evaluated using 20 repeated 80/20 train/test splits. Within each training partition (80%), elastic net, random forest, and XGBoost models were trained using either full dimensional or feature reduced predictors and three repeats of five-fold cross-validation. The evaluated data views included metabolomics only, taxonomy only, and the three concatenation variants. Out-of-fold (OOF) predictions from the metabolomics-only and taxonomy-only base models were generated within each repeat and combined with the outcome to train stacked models. OOF predictions were integrated using non-negative least squares (NNLS), simple averaging (AVG), elastic net (ENET), or partial least squares (PLS). Validation performance and feature importance were recorded from the cross-validation models, and final predictive performance was evaluated in the held-out 20% test partition. **(C)** Median top 20 feature-overlap summaries were generated for the metabolomics-only, taxonomy-only, and three concatenated data views. Pairwise comparisons evaluated feature-set agreement across elastic net, random forest, and XGBoost; transformed versus untransformed inputs; and full dimensional versus feature reduced inputs.

We next compared performance across single-omics and multi-omics integration strategies, focusing on two integration frameworks: concatenation and stacked generalization. In stacked generalization, predictions from individual base models are used as inputs to a final model. For concatenation, metabolomic and taxonomic features from each matched sample were combined into a single feature vector, and we evaluated three variants: unscaled concatenation, z-score-scaled concatenation, and z-score-scaled concatenation with block weighting (**Fig. 1A**). For stacked generalization, separate base models were first trained on the metabolomics and taxonomic abundance datasets, and their out-of-fold predictions were used to generate meta-features that were integrated using four strategies: simple averaging (AVG), weighted non-negative least squares (NNLS) [25], elastic net (ENET) [26], and partial least squares (PLS) [34] (**Fig. 1B**).

Each analysis incorporated grid search with 3 repeats of 5-fold cross-validation to optimize base-model hyperparameters, and model performance was evaluated using root mean squared error (RMSE) and mean absolute error (MAE) for continuous outcomes. Area under the receiver operating characteristic curve (AUROC) and area under the precision-recall curve (AUPRC) for binary outcomes. Uniform grid parameters were applied across analyses, and selected features, or, for stacked models, base-model feature importances, were extracted for downstream stability analyses. Finally, we compared predictive performance across single-omics, concatenation, and stacked integration strategies, and evaluated feature selection stability across machine learning models for single-omics and concatenated approaches (**Fig. 1C**).

### Competitive-set analysis identified model design choices that were consistently near-best

To compare model design choices across microbiome and metabolomics prediction tasks, we summarized performance using a competitive-set framework. For each dataset and performance metric, model configurations were ranked by their mean held-out test performance across 20 repeated train/test splits. For RMSE and MAE, configurations within 2% of the lowest mean error were retained (**Fig. 2A**). For AUROC and AUPRC, configurations within 2% of the highest mean performance were retained (**Fig. 3A**). Each dataset contributed equally to the aggregate summary, and when multiple configurations had identical mean performance values, the available credit for that value was divided equally among the tied configurations, with each receiving 1/*n* credit for *n* tied configurations. This approach accounted for cases in which preprocessing choices, such as scaling, did not affect performance. Because prediction endpoints differed in outcome scale and overall performance range, model configurations were compared within each dataset and metric rather than by directly pooling raw performance values across endpoints. The complete train and test performance summaries across repeated splits are provided in **Additional File 1**, and the underlying workbook containing the full performance tables, competitive-set memberships, and seed-level test differences is provided in **Additional File 2**. Together, these summaries supported identification of strategies that repeatedly ranked near the top across continuous and binary prediction tasks (**Fig. 2A**, **Fig. 3A**).

**Figure 2.**
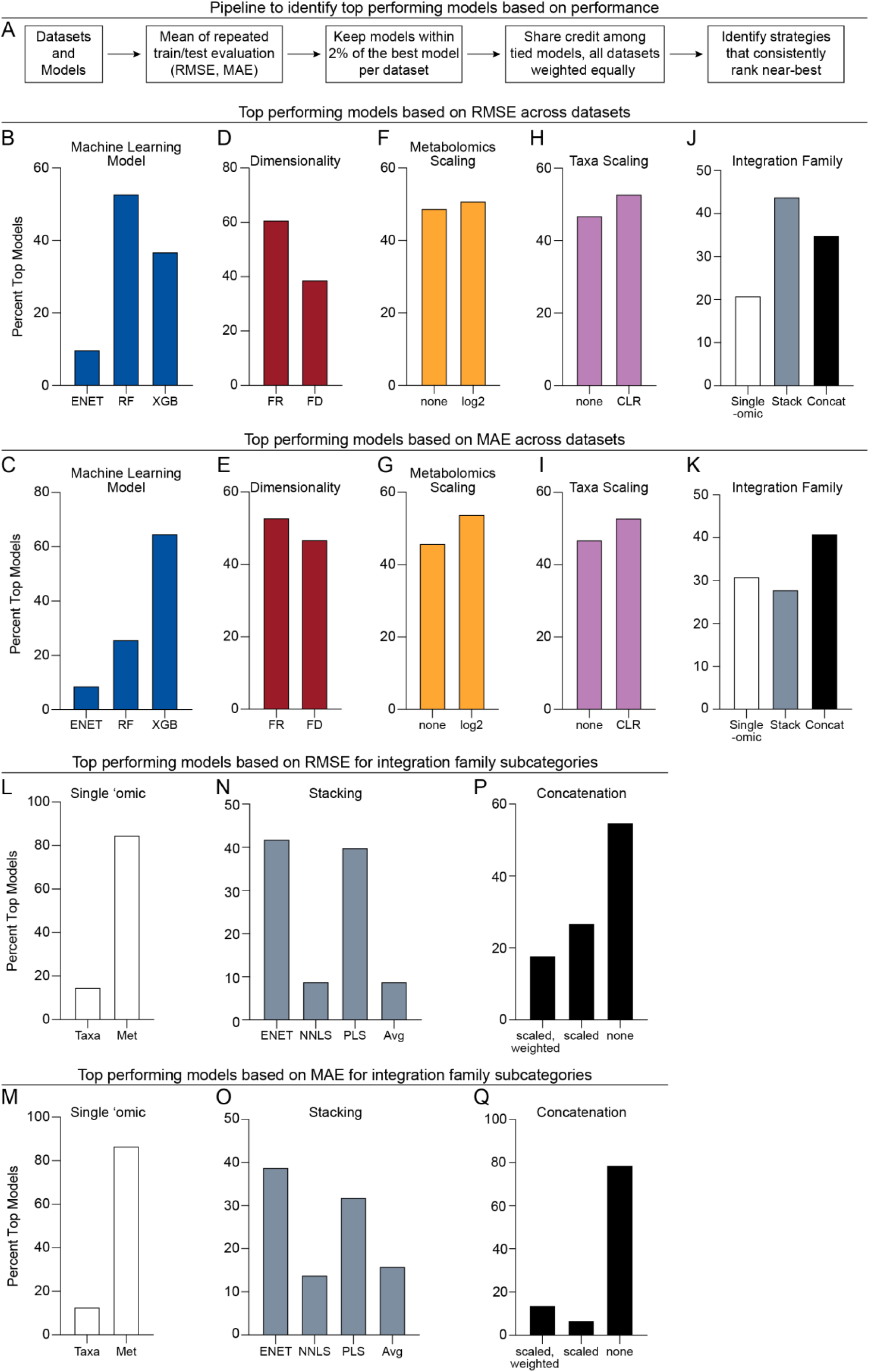
Continuous outcomes favored nonlinear learners, feature reduction, and metabolomics-dominant models. **(A)** Overview of the competitive-set analysis used to identify model configurations that ranked near-best across continuous prediction tasks. For each dataset and performance metric, held-out test performance was averaged across 20 repeated train/test splits. Configurations within 2% of the best-performing model were retained, with credit shared among tied models and datasets weighted equally. **(B–K)** Aggregate competitive-set summaries for continuous outcomes, shown separately for root mean squared error (RMSE; **B, D, F, H, J**) and mean absolute error (MAE; **C, E, G, I, K**). Panels summarize near-best model membership by machine learning algorithm (**B– C**), dimensionality setting (**D–E**), metabolomics scaling (**F–G**), taxonomic scaling (**H–I**), and integration family (**J–K**). **(L–Q)** Competitive-set summaries within integration-family subcategories for RMSE (**L, N, P**) and MAE (**M, O, Q**). Panels compare single-omic model type (**L–M**), stacked integration strategy (**N–O**), and concatenation variant (**P–Q**). Percentages represent dataset equal competitive-set credit among retained near-best configurations rather than pooled raw performance values. ENET, Elastic Net; RF, Random Forest; XGB, XGBoost; FR, feature reduced; FD, full dimensional; CLR, centered log-ratio; Met, metabolomics; Taxa, taxonomic relative abundance; Stack, stacked generalization; Concat, concatenation; NNLS, non-negative least squares; PLS, partial least squares; Avg, simple averaging.

**Figure 3.**
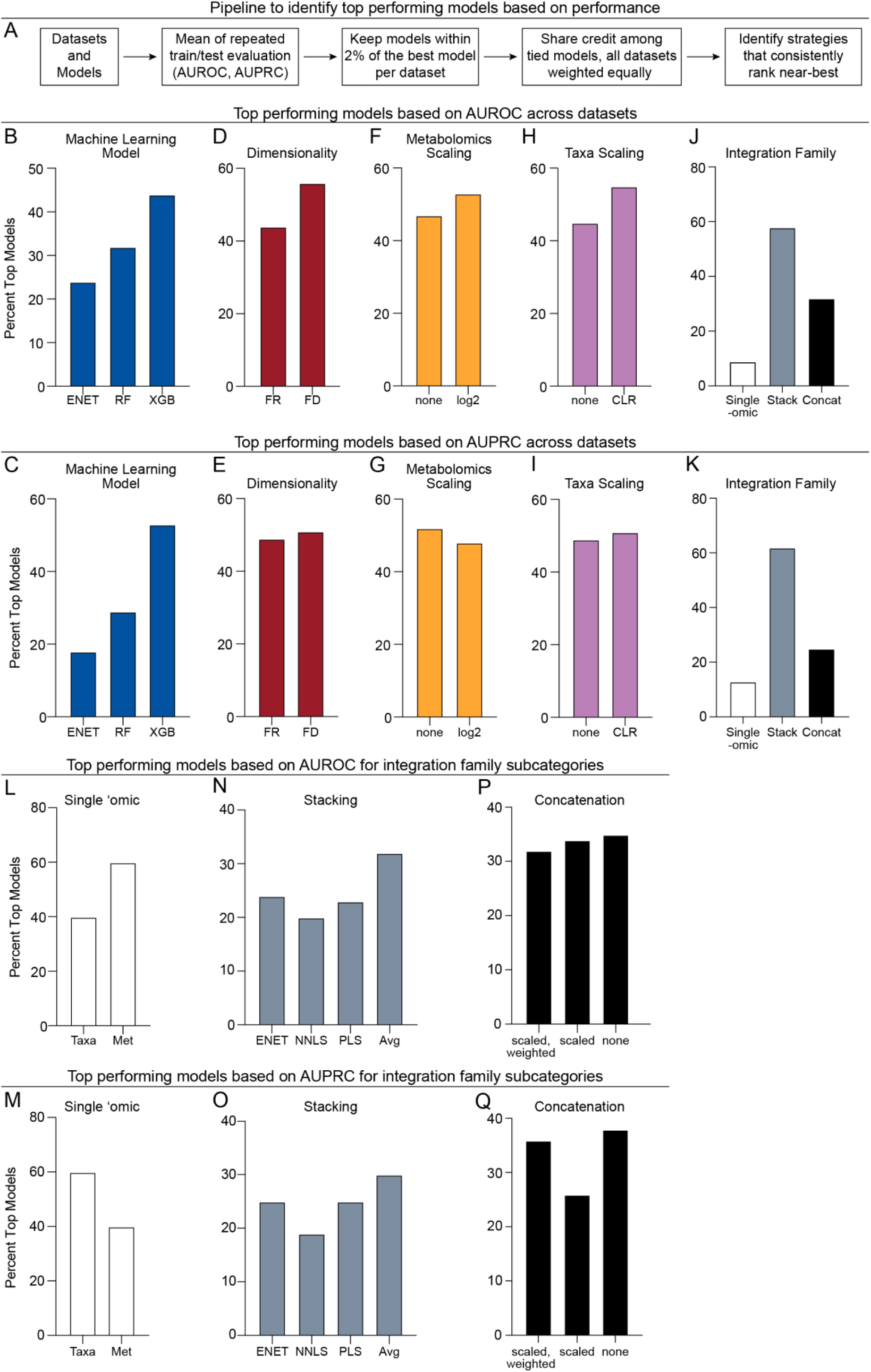
Binary classification outcomes favored stacked multi-omic models. **(A)** Overview of the competitive-set analysis used to identify model configurations that ranked near-best across binary classification tasks. For each dataset and performance metric, held-out test performance was averaged across 20 repeated train/test splits. Configurations within 2% of the best-performing model were retained, with tied configurations sharing credit and all datasets weighted equally. **(B–K)** Aggregate competitive-set summaries for binary outcomes, shown separately for area under the receiver operating characteristic curve (AUROC; **B, D, F, H, J**) and area under the precision-recall curve (AUPRC; **C, E, G, I, K**). Panels summarize near-best model membership by machine learning algorithm (**B–C**), dimensionality setting (**D–E**), metabolomics scaling (**F–G**), taxonomic scaling (**H–I**), and integration family (**J–K**). **(L–Q)** Competitive-set summaries within integration-family subcategories for AUROC (**L, N, P**) and AUPRC (**M, O, Q**). Panels compare single-omic model type (**L– M**), stacked integration strategy (**N–O**), and concatenation variant (**P–Q**). Percentages represent dataset-equal competitive-set credit among retained near-best configurations rather than pooled raw performance values. ENET, Elastic Net; RF, Random Forest; XGB, XGBoost; FR, feature reduced; FD, full dimensional; CLR, centered log-ratio; Met, metabolomics; Taxa, taxonomic relative abundance; Stack, stacked generalization; Concat, concatenation; NNLS, non-negative least squares; PLS, partial least squares; Avg, averaging; AUROC, area under the receiver operating characteristic curve; AUPRC, area under the precision-recall curve.

### Continuous outcomes favored nonlinear learners, feature reduction, and metabolomics-dominant models

Across the eight continuous prediction tasks, nonlinear ensemble methods made up most of the near-best models. The pattern was also clear across the competitive set for each dataset, where most of the best performing models were random forest or XGBoost models (**SFig. 1-2**). Random forest contributed the largest share of competitive-set credit for RMSE (53%), followed by XGBoost (37%) and elastic net (10%) (**Fig. 2B**). In contrast, XGBoost was dominant for MAE (65%), followed by random forest (26%) and elastic net (9%) (**Fig. 2C**). Feature reduced models were more frequently competitive than full dimensional models for both RMSE (61% vs. 39%) and MAE (53% vs. 47%), indicating that dimensionality reduction generally preserved near-minimum test error for continuous traits (**Fig. 2D-E**). Log_2_ transformation of metabolomics features and CLR transformation of taxa features provided modest advantages, but the magnitude of these effects was smaller than the effect of learner choice or integration strategy (**Fig. 2F-I**).

The best integration strategy for continuous outcomes depended on the error metric. Stacking was most often near-best for RMSE (44%), followed by concatenation (35%) and single-omic models (21%) (**Fig. 2J**). For MAE, concatenation had the largest share of near-best credit (41%), followed by single-omic models (31%) and stacking (28%) (**Fig. 2K**). Within single-omic models, metabolomics-only models accounted for most near-best single-omic performance for both RMSE (87%) and MAE (85%), whereas taxa-only models were less frequently represented among near-best single-omic models (**Fig. 2L-M**). Within stacking approaches, ENET and PLS stacking were the strongest contributors, while NNLS and simple averaging were less frequently near-best (**Fig. 2N-O**). Among concatenation approaches, unscaled concatenation was the dominant strategy, especially for MAE (79%) (**Fig. 2P-Q**). Dataset-level competitive-set maps for RMSE and MAE showed endpoint-specific variation in the exact retained configurations but supported the aggregate trends, with recurrent representation of metabolomics-only models, ENET or PLS stacking, and unscaled concatenation among near-best continuous-outcome models (**Figs. S1-S2**).

### Binary classification outcomes favored stacked multi-omic models

Across the nine binary classification tasks, XGBoost was the most consistently competitive learner, contributing 44% of near-best AUROC credit and 53% of near-best AUPRC credit (**Fig. 3B-C**). Random forest ranked second for both AUROC (32%) and AUPRC (29%), while elastic net contributed less often to the competitive set (24% for AUROC and 18% for AUPRC) (**Fig. 3B-C**). In contrast to the continuous outcomes, full dimensional models were more frequently competitive than feature reduced models for AUROC (56% vs. 44%) and were nearly balanced for AUPRC (51% vs. 49%) (**Fig. 3D-E**). AUROC was slightly higher for log_2_ transformed data, while AUPRC was slightly higher when the data was left untransformed (**Fig. 3F-G**). Both classification metrics were slightly higher when the taxa data were CLR transofrmed (**Fig. 3H-I**).

Multi-omic stacking was the clearest signal in the binary tasks. Stacked models accounted for the majority of near-best credit for both AUROC (58%) and AUPRC (62%), outperforming concatenation (32% and 25%) and single-omic models (9% and 13%), respectively (**Fig. 3J-K**). Among stacked models, simple averaging contributed the largest share of near-best credit for both AUROC and AUPRC, with ENET and PLS stacking also contributing substantially (**Fig. 3N-O**). In contrast, NNLS stacking was less frequently competitive (**Fig. 3N-O**). Among concatenation approaches, near-best credit was similar across variants for AUROC, with unscaled, scaled, and scaled-weighted concatenation contributing 35%, 34%, and 32%, respectively, and only modestly more variable for AUPRC, where unscaled and scaled-weighted concatenation contributed 38% and 36% compared with 26% for scaled concatenation, suggesting that concatenation variant was dataset dependent (**Fig. 3P-Q**). For single-omic classification models, the preferred modality depended on the metric: metabolomics-only models contributed more near-best credit for AUROC (60%), whereas taxa-only models contributed more for AUPRC (60%) (**Fig. 3L-M**). Dataset-level AUROC and AUPRC competitive-set maps similarly supported the aggregate binary-outcome pattern (**SFig. 3-4**). These show that stacked multi-omic models were repeatedly retained across classification endpoints, while concatenation and single-omic approaches appeared more selectively across datasets (**Figs. S3-S4**).

### Feature-stability analysis pipeline

We next wanted to evaluate whether alternative modeling and preprocessing choices identified the same predictive features. Feature importance was summarized across the 15 cross-validation fits within each of 20 outer runs, and features were ranked using a stability score that combined mean-importance percentile rank with outer-run presence frequency. The top 20 features were retained for each dataset, data view, model, and preprocessing condition, and pairwise overlap was summarized across the eight continuous or nine binary datasets. This produced 12 × 12 overlap matrices for the single-omics views and 24 × 24 matrices for the concatenated views (**Figs. 4A** and **5A**). Overlap ranged from 0%, indicating no shared features, to 100%, indicating identical top 20 feature sets. Throughout this section, transformed refers to log₂ transformation of metabolomic abundances and centered log-ratio (CLR) transformation of taxonomic abundances. A pseudocount of was applied before both transformations, where it was added to all metabolomic values and used to replace zero taxonomic abundances before CLR transformation. Feature reduced refers to limma-based reduction for metabolomics, Wilcoxon-based reduction for taxonomy, or both reductions for concatenated data.

**Figure 4.**
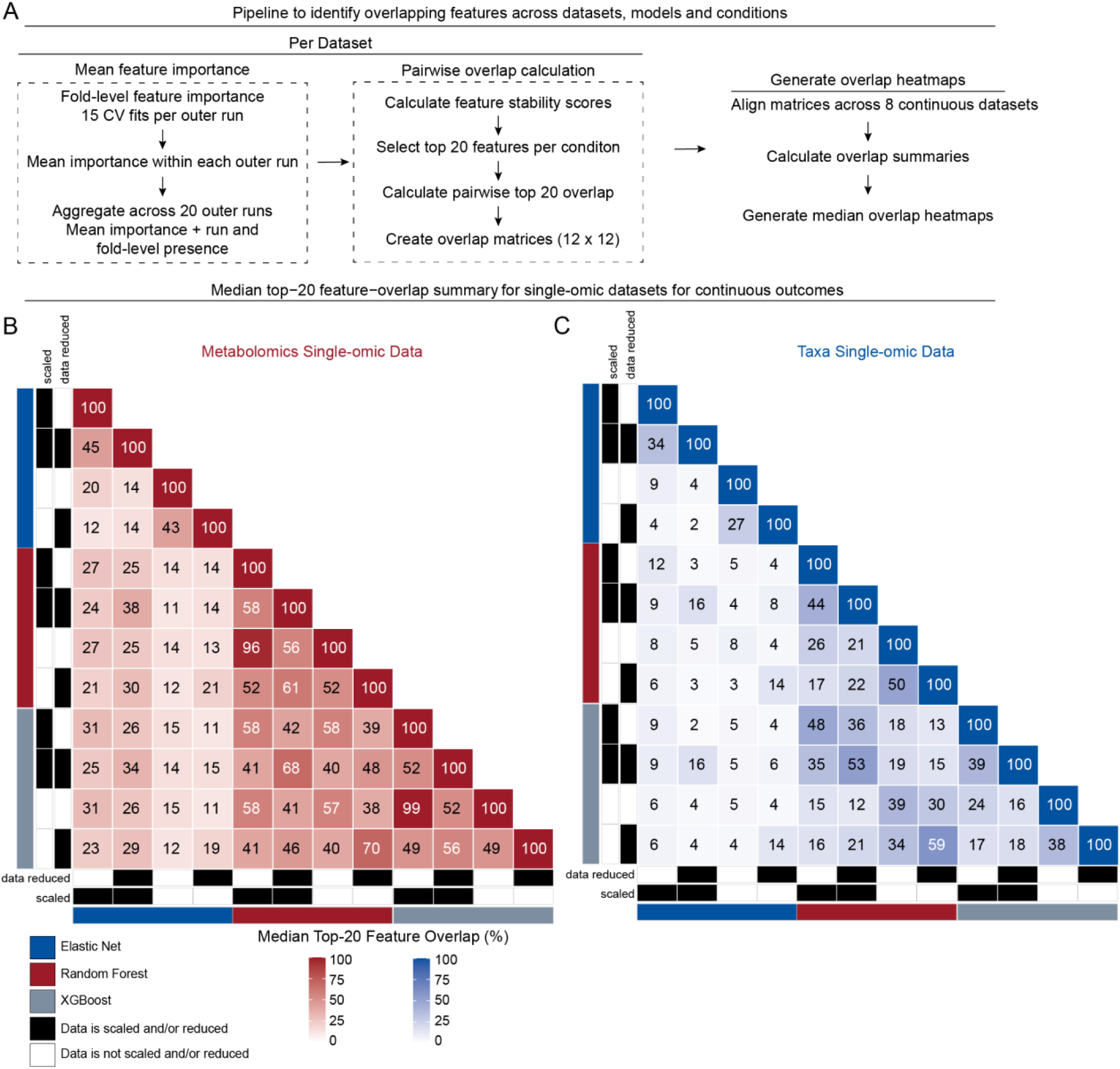
Metabolomics tree-based methods produce the most stable continuous feature rankings. **(A)** Workflow used to quantify feature selection stability and overlap. Feature importance was extracted for each dataset, data view, and pipeline condition from the 15 cross-validation fits within each of 20 outer runs, corresponding to three repeats of five-fold cross-validation per run. Fold-level importance values were averaged within each outer run and then across the outer runs in which the feature was present. Features absent from a fold or run were not assigned an importance of zero but contributed to the corresponding presence frequency. Mean importance was converted to a percentile rank, and the primary feature stability score was calculated as the importance percentile rank multiplied by outer-run presence frequency. A secondary score based on fold level presence frequency contributed to the prespecified tie-breaking procedure. Features were ranked by the primary stability score, and the 20 highest-ranking features were retained for each condition. Pairwise overlap was calculated (See Methods) producing a 12 × 12 overlap matrix for each single-omics view. Corresponding condition pairs were aligned across eight datasets with continuous outcomes, and overlap values were summarized across datasets. **(B-C)** Median top 20 feature-overlap matrices for the metabolomics and taxonomic single-omics views. Each row and column represents one of 12 pipeline conditions defined by three prediction models (elastic net, random forest, and XGBoost) and four view-specific preprocessing combinations. (**B**) Metabolomics conditions combined log_2_ transformation or no transformation with limma feature reduction or no reduction. (**C**) Taxonomic conditions combined centered log-ratio (CLR) transformation or no transformation with Wilcoxon-based feature reduction or no reduction. Blue, red, and gray annotation bars denote elastic net, random forest, and XGBoost, respectively. Black tiles in the preprocessing annotation tracks indicate that the corresponding transformation or feature reduction step was applied. White tiles indicate that it was not applied. Each heatmap cell reports the median percentage of shared top 20 features across the eight continuous-outcome datasets for the corresponding pair of conditions.

**Figure 5.**
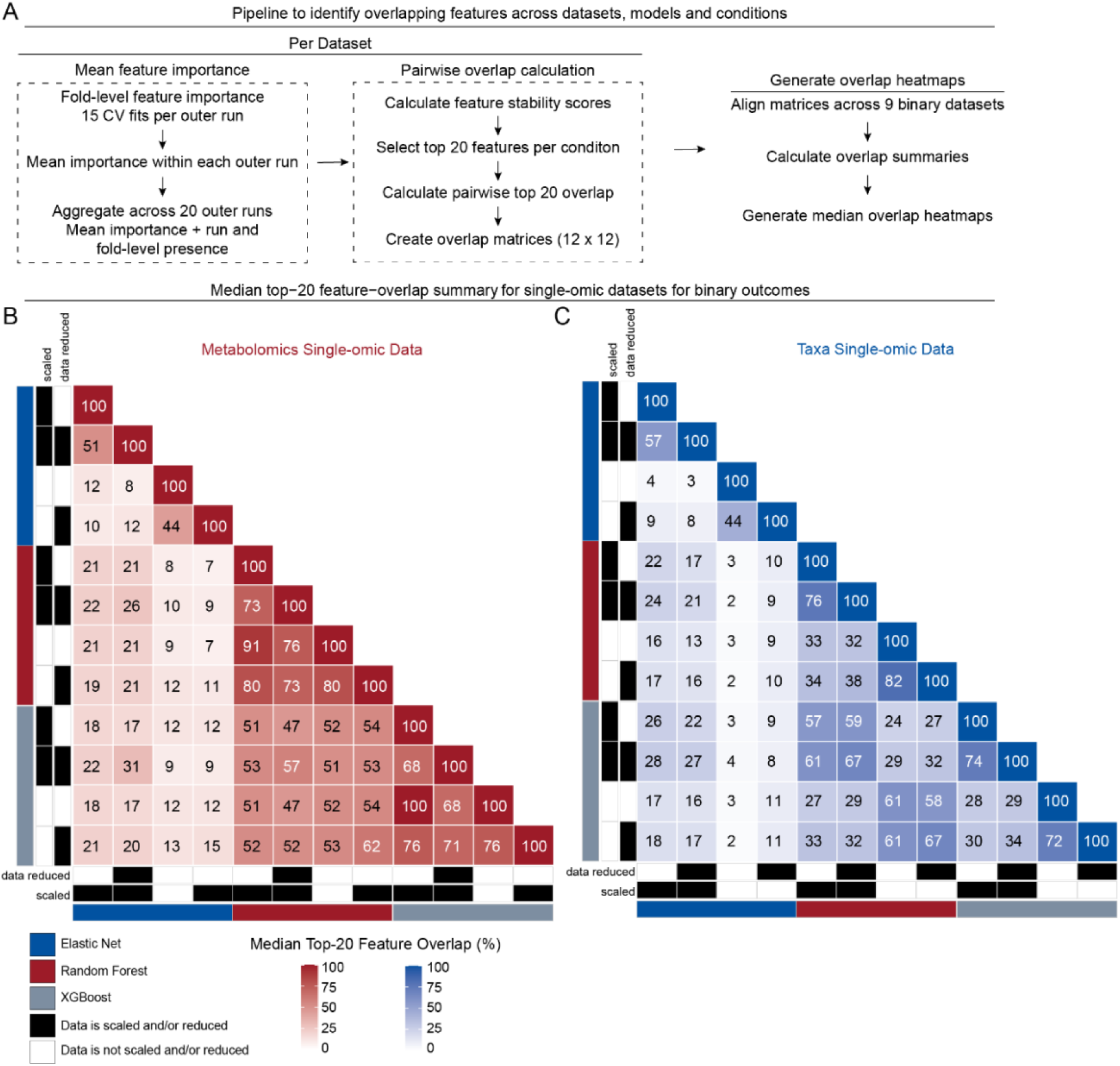
Nonlinear metabolomics feature rankings were highly reproducible for binary outcomes. **(A)** Workflow used to quantify feature-selection stability and overlap. For each study–outcome dataset, data view, and pipeline condition, feature importance was extracted from the 15 cross-validation fits within each of 20 outer runs, corresponding to three repeats of five-fold cross-validation per run. Fold-level importance values were averaged within each outer run and then across the outer runs in which the feature was present. Features absent from a fold or run were not assigned an importance of zero but contributed to the corresponding presence-frequency. Mean importance was converted to a percentile rank, and the primary feature stability score was calculated as the importance percentile rank multiplied by outer-run presence frequency. A secondary score based on fold level presence frequency contributed to the prespecified tie-breaking procedure. Features were ranked by the primary stability score, and the 20 highest-ranking features were retained for each condition Pairwise overlap was calculated (See Methods) producing a 12 × 12 overlap matrix for each single-omics view. Corresponding condition pairs were aligned across nine datasets with binary outcomes, and overlap values were summarized across datasets. **(B-C)** Median top 20 feature-overlap matrices for the metabolomics and taxonomic single-omics views. Each row and column represents one of 12 pipeline conditions defined by three prediction models (elastic net, random forest, and XGBoost) and four view-specific preprocessing combinations. (**B**) Metabolomics conditions combined log_2_ transformation or no transformation with limma feature reduction or no reduction. (**C**) Taxonomic conditions combined centered log-ratio transformation or no transformation with Wilcoxon-based feature reduction or no reduction. Blue, red, and gray annotation bars denote elastic net, random forest, and XGBoost, respectively. Black tiles in the preprocessing annotation tracks indicate that the corresponding transformation or feature reduction step was applied. White tiles indicate that it was not applied. Each heatmap cell reports the median percentage of shared top 20 features across the eight continuous-outcome datasets for the corresponding pair of conditions.

### Feature rankings for nonlinear models were most stable for metabolomics and most sensitive to taxonomic transformation

For continuous outcomes, full dimensional random forest and XGBoost metabolomics models were highly stable across transformed and untransformed data, retaining 96% and 99% of their top-ranked features, respectively. Agreement was lower after feature reduction, at 61% for random forest and 56% for XGBoost. Changing between full dimensional and feature reduced metabolomics produced approximately 49–58% overlap, indicating that feature reduction altered nonlinear model rankings more than metabolomics transformation. Elastic net was less stable overall and retained only 14–20% of its features when metabolomics transformation status changed (**Fig. 4B**).

Taxonomic rankings were less stable. Changing transformation status retained only 22–26% of random-forest features, 18–24% of XGBoost features, and 2–9% of elastic-net features. Switching between full dimensional and feature reduced taxonomic data had a similar impact, retaining 44–50% of random forest features and 38–39% of XGBoost features. Thus, metabolomics produced more reproducible continuous feature rankings than taxonomic data, and taxonomic transformation was the strongest source of single-omics feature selection instability (**Fig. 4C**).

Binary metabolomics rankings were highly reproducible for random forest and XGBoost. Changing transformation status in full dimensional data retained 91% of random forest features and 100% of XGBoost features. The corresponding feature reduced comparisons retained 73% and 71%. Changing between full dimensional and feature reduced metabolomics also retained 68–80% of nonlinear-model features. Elastic net was considerably more sensitive, retaining only 12% of its features when metabolomics transformation differed (**Fig. 5B**).

Taxonomic transformation again produced larger changes than feature reduction. Changing transformation status retained only 33–38% of random-forest features, 28–34% of XGBoost features, and 4–8% of elastic-net features. In contrast, changing between full dimensional and feature reduced taxonomy retained 76–82% of random-forest features and 72–74% of XGBoost features. Binary rankings were therefore generally more stable than continuous rankings, but metabolomics remained more reproducible than taxonomy (**Fig. 5C**).

### Continuous concatenated models were most sensitive to feature reduction

Across the three continuous concatenation variants, random forest and XGBoost were relatively robust to metabolomics transformation, retaining 80–95% and 68–100% of their features, respectively. Taxonomic transformation produced lower overlap of approximately 50–70%. The largest changes occurred between full dimensional and feature reduced data, which retained only 18–28% of random forest features and 8–12% of XGBoost features. Joint feature reduction was therefore the main source of instability in continuous concatenated models (**SFigs. S5–S7**).

Elastic net feature stability varied across the three concatenation strategies. When conditions differed only in taxonomic transformation, median overlap was 0–2% without post-concatenation z-score transformation or block weighting, increased to 28–48% after z-score transformation, and decreased to 8–28% when block weighting was added. Thus, z-score-transformed concatenation without block weighting yielded the most reproducible elastic net top 20 feature sets, whereas random forest and XGBoost showed similar feature-set stability across all three concatenation variants (**SFigs. S5–S7**). The greater overlap seen after transforming the taxonomic data in the concatenated models may partly be due to differences in the metabolomics features. It does not necessarily mean that the taxonomic features themselves were more stable.

### Binary nonlinear concatenated models were the most stable across preprocessing variants

Binary concatenated random forest and XGBoost models were more robust than their continuous counterparts (**SFigs. 8-10**). Changing metabolomics transformation retained 80–100% of their top-ranked features, changing taxonomic transformation retained 40–75%, and changing between full dimensional and feature reduced data retained 60–85%. The difference between binary and continuous models was greatest for feature reduction, where binary nonlinear models retained 60–85% of their features, compared with only 8–28% for continuous models (**SFigs. 8–10**).

As in the continuous analysis, post-concatenation z-score transformation most clearly improved the reproducibility of elastic-net top-20 feature sets. Median overlap increased from 0–10% to 25–45% for conditions differing only in taxonomic transformation and from 30–45% to 45–70% for conditions differing only in dimensionality (**SFigs. S8–S9**). Adding block weighting reduced these ranges to 20–35% and 15–40%, respectively. Random forest and XGBoost remained similarly stable across all three concatenation variants (**SFig. S10**).

### Full dimensional metabolomics with nonlinear learners produced the most stable feature rankings

Across all five data views, the most stable feature rankings were produced by full dimensional metabolomics models using random forest or XGBoost (**Figs. 4-5**, **SFigs. 5-10**). When metabolomics transformation status changed, these models retained 96–99% of their features for continuous outcomes and 91– 100% for binary outcomes. Taxa-only models were the least stable across transformation, retaining only 18– 26% of nonlinear model features for continuous outcomes and 28–38% for binary outcomes. Concatenated views generally fell between these extremes, although continuous concatenated models were particularly sensitive to feature reduction. Overall, 1) random forest and XGBoost were more stable than elastic net; 2) binary rankings were more stable than continuous rankings; and 3) z-score-transformed concatenation without block weighting provided the clearest stability benefit only for elastic net.

## Discussion

This study systematically compared predictive performance and feature selection stability across four matched microbiome–metabolomics datasets using eight continuous outcomes, and nine binary outcomes. The analysis evaluated three learners, omics-specific transformations, full dimensional and feature reduced inputs, and multiple single and multi-omics integration strategies (**Fig. 1A–B**). Feature stability was then assessed by comparing the top 20 feature sets selected across learners and preprocessing conditions (**Fig. 1C**). The competitive-set framework emphasized configurations that were repeatedly near-best across datasets and metrics, rather than overinterpreting small differences among closely performing models. Overall, nonlinear learners were more consistently competitive than elastic net, but the preferred integration strategy depended on the outcome type and performance metric.

For continuous outcomes, random forest contributed most often to near-best RMSE performance, whereas XGBoost contributed most often to near-best MAE performance (**Fig. 2B-C**, **SFigs. 1-2**). Feature reduced models were more frequently competitive than full-dimensional models, indicating that substantial dimensionality reduction often preserved predictive performance (**Fig. 2D-E**). Among single-omics models, metabolomics accounted for 85-87% of near-best credit (**Fig. 2L-M**), which may reflect closer links between the evaluated clinical outcomes and host or microbial metabolism. Integration performance was metric-dependent: stacking was most often near-best for RMSE, whereas concatenation was favored for MAE (**Fig. 2J-K**). ENET and PLS were the strongest stacking methods, while concatenation without post-concatenation z-scoring or block weighting was most competitive, particularly for MAE (**Fig. 2N-O** and **Fig. 2P-Q**).

Binary outcomes showed a clearer benefit from prediction-level integration. XGBoost was most consistently competitive for both AUROC and AUPRC, and stacked models accounted for 58% and 62% of near-best credit, respectively (**Fig. 3B-C**, **Fig. 3J-K**, **SFigs. S3-S4**). Simple averaging was the strongest stacking method, suggesting that complementary omics signals could be combined without a complex meta-learner; ENET and PLS also accounted for substantial shares of near-best performance credit (**Fig. 3N-O**). This finding contrasts with a previous study in which Bayesian additive regression tree (BART)-NNLS outperformed alternative integration approaches [16]. The discrepancy may reflect differences in base-learner and meta-learner configurations or evaluation settings, particularly because BART performed well for individual-layer prediction in that study but was not evaluated here. Single-omics performance was also metric-dependent. Metabolomics was more often competitive for AUROC, whereas taxonomy was more often competitive for AUPRC (**Fig. 3L-M**), underscoring the importance of evaluating both metrics, especially for imbalanced outcomes.

Preprocessing effects were generally smaller than learner and integration effects, and no transformation or dimensionality setting was uniformly preferred. Feature reduction was more often near-best for continuous outcomes, whereas full dimensional models were modestly favored for binary AUROC and nearly balanced with feature reduced models for AUPRC (**Figs. 2D-E and 3D-E**). Transformations of metabolomic and taxonomic data produced only modest predictive differences (**Figs. 2F-I and 3F-I**). However, prior benchmarking found that microbial-load-aware quantitative approaches outperformed computational strategies for diversity and association analyses [35]. Together, these findings support selecting preprocessing methods by dataset and endpoint, particularly because similar predictive performance can mask substantial differences in selected features.

Full dimensional metabolomics models using random forest or XGBoost yielded the most reproducible top 20 feature sets (**Figs. 4A and 5A**). Across metabolomic transformation conditions, these models retained 96-99% of features for continuous outcomes and 91-100% for binary outcomes (**Figs. 4B and 5B**). Across taxonomic transformation conditions, nonlinear models retained only 18-26% of features for continuous outcomes and 28-38% for binary outcomes (**Figs. 4C and 5C**). Elastic net was even more sensitive to preprocessing, retaining fewer than 10% of selected features in some taxonomic comparisons. This pattern is consistent with learner structure: elastic net emphasizes sparse linear associations, whereas random forest and XGBoost capture nonlinear effects and interactions across distributed predictors [36]. Additionally, binary feature sets were generally more reproducible than continuous feature sets, particularly when dimensionality changed.

Among concatenated models, continuous random forest and XGBoost were relatively stable across metabolomics transformation but retained only 8-28% of features when full dimensional and feature reduced inputs were compared (**SFigs. S5-S7**). This sensitivity may partly reflect the larger metabolomics feature block, because reducing a larger candidate pool can substantially change the composition of the combined top 20 set. Binary concatenated models were more robust, retaining 60-85% of nonlinear-model features across dimensionality settings (**SFigs. S8-S10**). Post-concatenation z-score transformation without block weighting most clearly improved elastic net feature agreement, while random forest and XGBoost showed no consistent stability advantage for any concatenation variant. Because the most stable concatenation variant was not uniformly the best-performing one (**Figs. 2P-Q and 3P-Q**), predictive accuracy alone is insufficient when biological interpretation is also a goal.

These findings align with prior microbiome studies showing that transformation and model choice can alter feature importance despite relatively stable classification accuracy [37] and that microbial feature-selection workflows can be unstable [38]. In our analyses, strong predictive performance did not ensure agreement on the same biological features, and different learners identified distinct but similarly predictive representations. Features intended for mechanistic interpretation or candidate prioritization should therefore be evaluated across resampling runs, preprocessing choices, and plausible model classes rather than selected from a single fitted model.

Several limitations should be considered. The analysis included four cohorts and 17 outcomes, but multiple endpoints were derived from the same studies and therefore do not represent 17 independent cohorts. Validation in additional diseases, populations, and external datasets will be necessary to determine how broadly the observed performance and stability patterns generalize. Taxonomic profiles were generated using Kraken and Bracken, and alternative profiling tools or taxonomic resolutions could change both predictive performance and selected features. We also evaluated only one feature reduction approach for each omics type, so the observed effects may partly reflect these specific methods. Prior metabolomics benchmarking has shown that feature selection and feature extraction choices can alter classification performance [39], limiting generalization to dimensionality reduction methods not evaluated here. Unequal feature-block sizes may further influence concatenated top 20 sets, making combined overlap an imperfect measure of the stability of either omics block individually.

Overall, the results do not support a single default strategy for microbiome-metabolomics prediction. Nonlinear learners and metabolomics-dominant models were most consistently competitive for continuous outcomes (**Fig. 2**), whereas stacked multi-omics models provided the clearest benefit for binary classification (**Fig. 3**). Full dimensional metabolomics models using random forest or XGBoost produced the most reproducible feature sets, while taxonomic transformation and feature reduction in continuous concatenated models caused the largest shifts in selected features (**Figs. 4-5**; **SFigs. S5-S10**). Table 2 summarizes practical recommendations for microbiome-metabolomics machine learning (**Table 2**). Future work should extend this framework to additional cohorts, profiling and feature reduction methods, refined hyperparameter searches, and alternative integration approaches such as DIABLO [40] and MOFA+ [41]. Most importantly, multi-omics machine learning studies should report predictive performance and feature selection stability together, particularly when model outputs inform biological interpretation or feature prioritization.

**Table 2.**
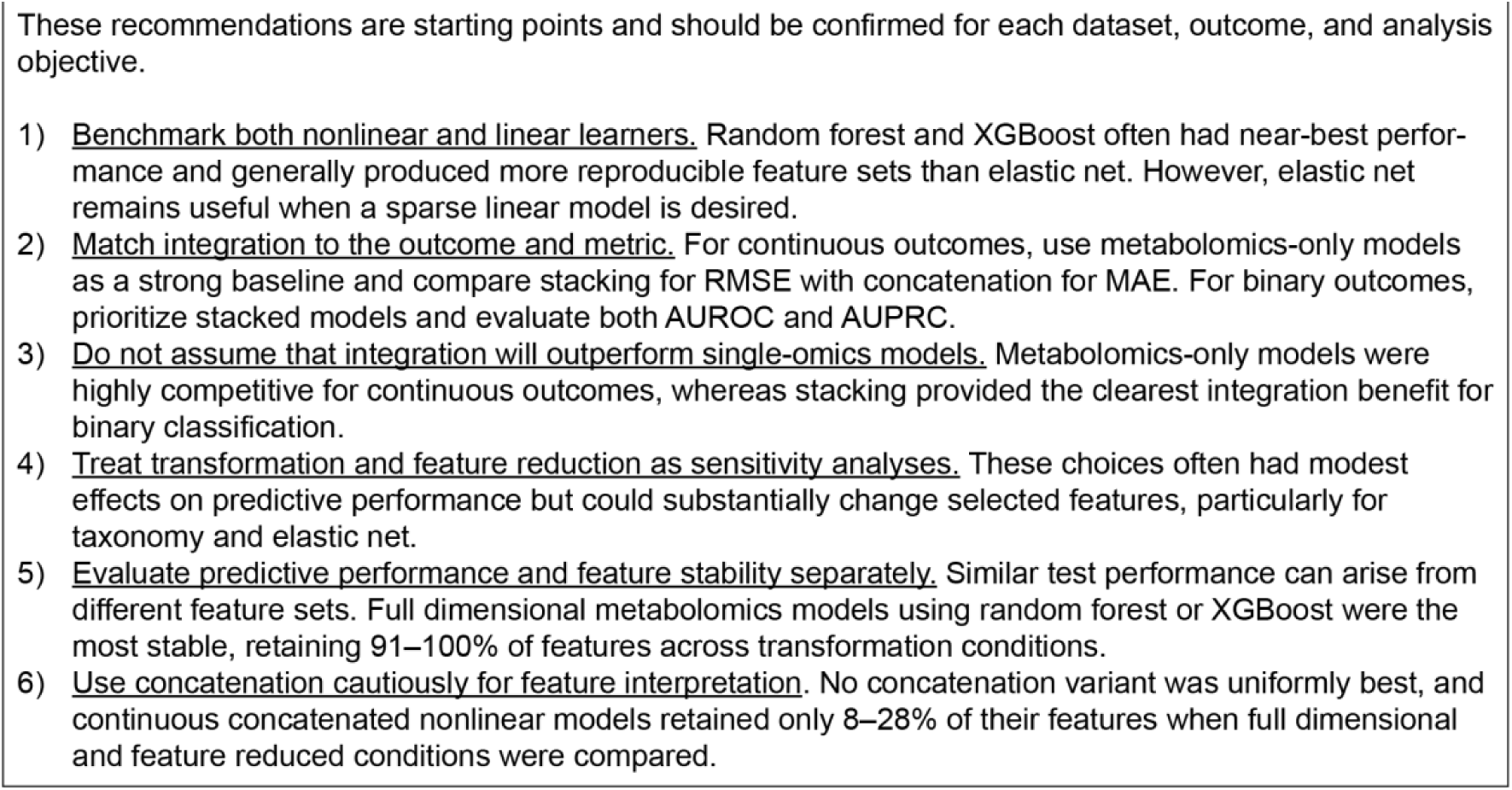
Practical recommendations for microbiome-metabolomics machine learning.

## Methods

### Data sources and preprocessing

Data from Franzosa *et al.* (2019) [11], Yachida *et al.* (2019) [14], Erawijantari *et al.* (2020) [23], and Wang *et al.* (2020) [24] were obtained as paired taxonomic relative-abundance tables, metabolomics, and metadata tables from Müller *et al.* (2022) [42]. Briefly, raw metagenomic shotgun sequencing FASTQ files from the original studies were processed with fastp v.0.23.2 [43] for quality filtering, adapter trimming, and deduplication, followed by host-read removal with Bowtie v.2.3.5 [44] against the human reference genome (Genome Reference Consortium Build 38). Taxonomic classification and relative-abundance estimation were performed with Kraken v.2.1.1 and Bracken v.2.8 using GTDB v.207 as the reference database [45,46]. Samples with fewer than 50,000 reads were excluded. For the present analyses, genus and species-level taxonomic data were obtained from genera.tsv and species.tsv files. Metabolomics data were obtained from mtb.tsv with metabolite identifiers as originally reported, and metadata were obtained from metadata.tsv [42].

For integrative modeling, genus and species-level taxonomic tables were concatenated within each dataset. In the modeling input, metabolite features and taxonomic features were identified by the prefixes m and t. Rows containing missing values in the response variable, metabolite features, taxonomic features, or stratification variable were removed prior to analysis.

### Software environment

Machine learning analyses were implemented in R using caret v.7.0-1 [47], with elastic-net models fit using glmnet v.4.1-10 [26], random forests using randomForest v.4.7-1.2 [48], extreme gradient boosting using xgboost v.3.2.0 [49], partial least-squares models using pls v.2.9-0 [34], kernel-based models using kernlab v.0.9-33 [50], and non-negative least-squares fitting using nnls v.1.6 [25]. Model performance was evaluated using pROC v.1.19.0.1[51], PRROC v.1.4 [52], and MLmetrics v.1.1.3, and agreement-related analyses were performed using irr v.0.84.1. The package tidyverse v.2.0.0 [53], and argparse v.2.3.1 were used. Differential metabolite filtering was performed using limma v.3.64.1 [54].

### Study design and resampling

We developed a multi-view learning framework to evaluate metabolomics-only, taxonomy-only, and integrated metabolomics and taxonomy models for binary and continuous outcomes. For each study, data were partitioned into 80% training and 20% held-out test sets using caret::createDataPartition, and model development within each outer training set used 3 repeats of 5-fold cross-validation. To assess robustness to data partitioning, the entire train/test workflow was repeated using 20 seeds (1001-1020).

### Feature transformation and reduction

Before each outer train/test partition was generated, taxonomic features with a summed abundance below 1 × 10^−4^across the complete input dataset were excluded. Within each cross-validation fold, the specified metabolomics and taxonomic transformations were applied separately to the training and validation subsets. Feature selection was performed using only the fold-specific training data, and the selected feature sets were applied unchanged to the corresponding validation data.

Metabolite abundances were either left untransformed or log_2_ transformed. Taxonomic abundances were either left untransformed or subjected to a centered log-ratio (CLR) transformation. Zero taxon abundances were replaced with a pseudocount of 1 × 10^−9^. CLR values were then calculated using the natural logarithm by subtracting the within-sample mean log abundance from the log abundance of each taxon.

Feature reduction was performed separately for the metabolomics and taxonomic blocks using only the training samples from each cross-validation fold. Metabolites were evaluated using empirical Bayes–moderated linear models implemented in limma, with the comparison defined by the two levels of the configured binary stratification variable. Metabolites were ranked by their unadjusted *P* values. Up to 200 metabolites with the smallest *P* values were retained.

Taxa were evaluated individually using two-sided Wilcoxon rank-sum tests comparing the two levels of the binary stratification variable. The *P* values were adjusted using the Benjamini–Hochberg false discovery rate procedure, and taxa with adjusted *P* ≤ 0.05 were retained. If no taxa met this threshold, up to 200 taxa with the smallest unadjusted *P* values were selected. If at least one but fewer than 10 taxa met the threshold, the selected set was supplemented with the next most significant taxa until 10 taxa were retained.

The unscaled concatenated representation contained the same selected features and the same transformed values as the corresponding metabolomics-only and taxonomy-only representations for samples shared across those datasets. The two additional concatenated representations (described below) were generated from this same selected feature set. Following cross-validation, the block-specific transformation and feature-selection procedures were repeated using the complete training set. The resulting metabolite and taxon feature sets were applied to the held-out test set, and the concatenated test representations were constructed from the union of these selected features.

### Integration strategies

We evaluated direct concatenation and stacked generalization. For direct concatenation, the metabolite and taxonomic features retained after fold-specific selection were combined column-wise:

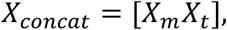

where *X_m_* and *X_t_* denote the metabolomics and taxonomic predictor matrices. Each matrix contained the same *n* samples and *P_m_* or *P_t_* retained features, such that *X_concat_* contained *P_m_* + *P_t_* predictors. Three concatenated variants were tested: unscaled concatenation, z-score normalization (referred to as z-score scaling) of all predictors, and z-score scaling followed by block weighting. For block weighting, metabolite and taxonomic features were multiplied by 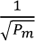 and 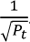.

For stacked integration, separate base learners were trained using the metabolomics and taxonomic views. Within each outer training partition, 3 repeats of 5-fold cross-validation was used to generate cross-fitted predictions. In each repeat, every training sample was predicted only by base learners fitted without that sample. For the metabolomics and taxonomic views, the out-of-fold predictions obtained for each samples across repeats were averaged, yielding one metabolomics prediction and one taxonomic prediction per outer training sample. The concatenated representations were evaluated as separate direct concatenation models and were not used as meta features. These averaged out-of-fold predictions, and no predictions from base models fitted to the complete outer training partition, were used as predictors for the second-stage meta-models.

Four prediction-level integration strategies were evaluated. Equal-weight averaging combined the two base-model predictions according to

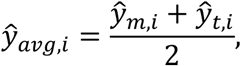

where ŷ*_m_*_,*i*_ and ŷ*_t_*_,*i*_, denote the metabolomics and taxonomic predictions for sample *i*. Equal weight averaging was a fixed combination rule and did not require fitting a meta-learner. For non-negative least squares (NNLS) integration,

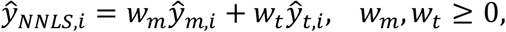

where *w_m_* and *w_t_* are non-negative weights estimated from the averaged out-of-fold metabolomics and taxonomic predictions. The NNLS model was fitted without an intercept, and the weights were not constrained to sum to one. Elastic net integration used the two averaged out-of-fold predictions as predictors in a regularized meta-model. Values of *α* from 0 to 1 in increments of 0.1 and *λ* from 10^2^ to 10^-2^ were evaluated using 5-fold cross-validation, after which the selected model was refitted using all outer training out-of-fold predictions. PLS integration similarly used the two averaged out-of-fold predictions as predictors, with the number of components selected by minimizing cross-validated PRESS before refitting the model using all outer-training out-of-fold predictions.

### Base learners

Elastic net, random forest, and extreme gradient boosting (XGBoost) were selected as base learners. Elastic net models were fit in caret using glmnet. Elastic net combines *L*_1_and *L*_2_ penalties, with the mixing parameter *α* controlling the relative contribution of the two penalties and *λ* controlling the overall degree of regularization. Tuning evaluated *α* values from 0 to 1 in increments of 0.05 and *λ* values from 10^-4^ to 10^1^.

Random forest models were fit in caret using randomForest, with candidate mtry values ranging from 1 to √*p*, where *p* denotes the number of predictors. For elastic net and random forest models, hyperparameters were selected by maximizing the area under the receiver operating characteristic curve (ROC AUC), calculated using pROC, for binary outcomes and by minimizing root-mean-square error (RMSE) for continuous outcomes.

XGBoost models were fit using a custom wrapper around xgboost. The default tuning grid included nrounds values of 300 and 600, max_depth values of 3 and 6, eta values of 0.05 and 0.10, colsample_bytree values of 0.7 and 1.0, and subsample values of 0.7 and 1.0. Binary models used the binary:logistic objective and were tuned by maximizing ROC AUC, whereas continuous models used the reg:squarederror objective and were tuned by minimizing RMSE. Histogram-based tree construction was used.

### Model fitting, prediction, and evaluation

Within each inner cross-validation fold of an outer training partition, each base learner was trained independently on the metabolomics-only, taxonomy-only, and concatenated feature sets. Predictions were generated only for the corresponding held-out validation fold, such that every out-of-fold prediction was produced by a base model that had not been trained on that sample. For binary elastic net and random forest models, predicted outputs were positive-class probabilities from caret::predict.train(…, type = “prob”). For binary XGBoost models, predicted outputs were logistic probabilities from the trained booster. For the data view, out-of-fold predictions were averaged across repeats for each sample.

The averaged out-of-fold predictions from the metabolomics-only and taxonomy-only base models served as meta-features for the second-stage integration models. NNLS integration was fitted using nnls. Elastic net integration used glmnet with *α* values from 0 to 1 in increments of 0.1 and *λ* values from 10^2^ to 10^-2^. PLS integration used the pls package, with the number of components selected by minimizing the cross-validated predicted residual error sum of squares (PRESS).

Using the transformations and feature sets re-estimated from the complete outer training set, final base learners were fitted for the metabolomics-only, taxonomy-only, and each concatenated representation. The final concatenated models directly generated predictions for the corresponding held-out outer test representations. For stacked integration, the final metabolomics and taxonomic base learners, rather than the fold-specific cross-validation models, were used to generate the two predictions for each held-out outer test sample. These predictions were supplied to the fixed equal-weight averaging rule or to the NNLS, elastic-net, and PLS meta-models previously fitted using the outer-training out-of-fold predictions. The meta-models were not refitted using outer-test data, and the outer-test outcomes were used only to calculate performance metrics. Binary performance was summarized using area under the receiver operating characteristic (AUROC), area under the precision-recall curve (AUPRC), sensitivity, specificity, precision, F1 score, accuracy, and balanced accuracy. Continuous performance was summarized using root mean square error (RMSE) and mean absolute error (MAE).

### Performance summarization and competitive-set analysis

Endpoint-level performance outputs from the repeated train/test workflow were post-processed to generate the aggregate summaries shown in Figs. 2 and 3 and the corresponding Figs. S1-S4 and Additional Files 1 and 2. For each dataset, model category, pipeline configuration, split, and performance metric, seed-level estimates from the 20 repeated train/test partitions were summarized as the mean across seeds.

To identify model-design choices that were consistently near-best across endpoints, we performed a competitive-set analysis using held-out test performance only. Continuous endpoints were summarized using RMSE and MAE. Binary endpoints were summarized using AUROC and AUPRC. Within each dataset and metric, model configurations were ranked by their mean test performance across the 20 train/test splits. For RMSE and MAE, the best-performing configuration was defined as the model with the lowest mean error, and all configurations with mean error less than or equal to 1.02 times this value were retained. For AUROC and AUPRC, the best-performing configuration was defined as the model with the highest mean performance, and all configurations with mean performance greater than or equal to 0.98 times this value were retained. For each retained configuration, the absolute and proportional difference from the best-performing configuration was recorded.

Pipeline identifiers were parsed to assign each retained configuration to the model factors described above. Each dataset contributed one unit of credit per metric to prevent endpoints with larger competitive sets from dominating the aggregate summaries. In the dataset-equal summaries used for **Figs. 2 and 3**, this credit was divided equally among all configurations retained for that dataset and metric. Exact ties in mean test performance were also identified; in the exported raw competitive-set tables, configurations with identical mean performance values shared credit for that performance value, with each tied configuration receiving 1/*n* credit for *n* tied configurations. This tie-sharing procedure was retained to document cases in which preprocessing choices, such as scaling, produced identical mean performance.

Competitive-set credits were then aggregated across datasets separately for each metric and model factor. Percentages were calculated by dividing the summed credit for each factor level by the total credit within that factor and metric. For integration subcategories, percentages were calculated within the relevant integration family. Dataset-level competitive-set maps were generated from the same retained configurations to show which model designs entered the competitive set for each endpoint and metric. The complete tabular outputs, including train/test performance matrices, competitive-set membership tables, frequency summaries, and seed-level test summaries, are provided in Additional File 2.

### Feature importance stability and overlap analysis

Feature importance was extracted from the 15 cross-validation models fitted within each outer run (3 repeats × 5 folds). For elastic net, feature importance was defined as the absolute value of the fitted coefficient at the selected regularization parameter, excluding the intercept. Random-forest importance was obtained using caret::varImp, and XGBoost importance was defined as gain obtained using xgb.importance.

Feature importance was summarized separately for each study–outcome dataset, pipeline condition, and data view. The evaluated views comprised metabolomics alone, taxonomy alone, and three concatenated variants. Each single-omics view included 12 prespecified model–preprocessing conditions, corresponding to three prediction models crossed with four view-specific preprocessing combinations. For the metabolomics view, log_2_ transformation or no transformation was combined with either no feature reduction or limma feature reduction. For the taxonomic view, CLR transformation or no transformation was combined with either no feature reduction or Wilcoxon-based feature reduction. Preprocessing settings for the omitted omics block were not treated as separate conditions because they did not affect feature selection within the single-omics view. Each concatenated view included 24 prespecified model preprocessing conditions, jointly defined by the prediction model and the metabolomics and taxonomic transformation and feature reduction settings. Within each of the 20 outer runs, the mean importance of each feature was calculated. These run-level means were then averaged across the outer runs when the feature was present. Features absent from a fold or run contributed to the presence-frequency denominator but were not assigned an importance of zero when calculating conditional mean importance.

Two measures of feature selection frequency were calculated. Outer-run presence frequency was defined as the number of outer runs in which a feature was assigned an importance value divided by the 20 total runs. Fold-level presence frequency was defined as the number of cross-validation fits in which a feature was assigned an importance value divided by the total number of fold-level fits across runs, corresponding to 300 expected fits per pipeline condition (20 runs × 15 fits). Because the numerical scales of feature-importance measures differed among modeling algorithms, mean importance was converted to a percentile rank ranging from 0 to 1 within each dataset, pipeline condition, view, and importance stratum. A primary feature-stability score was then calculated as:

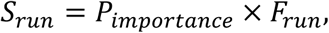

where *P_importance_* is the percentile rank of mean importance and *F_run_* is the outer-run presence frequency. A secondary fold-level stability score was calculated as:

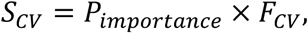

where *F_CV_* is the fold-level presence frequency.

Within each pipeline condition and data view, features were ranked by the primary stability score, and the 20 highest-ranking features were retained. Ties were resolved sequentially using outer-run presence frequency, the fold-level stability score, mean-importance percentile rank, and feature name. The top-20 overlap (%) for each dataset is found in Additional File 3. Feature-set agreement was evaluated separately within each data view. For each pair of pipeline conditions (12 for single-omics and 24 for concatenation), top 20 overlap was calculated as

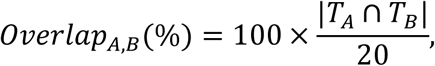

where *T_A_* and *T_B_* denote the top 20 feature sets for conditions A and B. An overlap of 100% indicated identical top 20 feature sets, whereas an overlap of 0% indicates that no selected features were shared. For each study– outcome dataset, five view-specific 12×12 (single-omics) or 24×24 (concatenation) overlap matrices were generated, corresponding to the metabolomics, taxonomy, unscaled concatenated, scaled concatenated, and scaled-and-weighted concatenated views.

To assess the reproducibility of preprocessing and modeling effects across datasets, overlap values were summarized separately for binary and continuous endpoints. The binary analysis included nine study–outcome datasets, and the continuous analysis included eight study–outcome datasets. For every within-view pair of pipeline conditions, the mean, median, standard deviation, and interquartile range of the top 20 overlap percentage were calculated across datasets in the corresponding endpoint class. Median 12×12 (single-omics) or 24×24 (concatenation) heatmaps were generated separately for each of the five data views and for binary and continuous endpoints.

### Split diagnostics and reproducibility

To quantify reuse of samples across repeated outer train/test partitions, test indices from all 20 runs were recorded and pairwise overlap were computed. The pipeline generated structured outputs including run documentation, fold-level and repeat-level performance tables, feature-importance files, feature-count logs, stability summaries, and serialized RData files. Reproducibility was further supported by explicit seed setting at the outer split, repeat/fold generation, learner-specific internal resampling, and final model-fitting stages.

### Use of artificial intelligence tools

The authors used generative AI tools for limited assistance with code development and manuscript wording. The study ideas, design, analyses, interpretation, and conclusions were developed by the authors. All AI-assisted code and text were independently checked, revised, and approved by the authors, who accept full responsibility for the final content. This statement is included to ensure transparent disclosure of AI use.

## Declarations

**Ethics Approval and Consent to Participate** – Not applicable

**Consent for Publication** – Not applicable

## Availability of Data and Material

Data is free and available through: https://github.com/borenstein-lab/microbiome-metabolome-curated-data/ [42]. Code for this integrative analysis and associated analysis are free and available through: https://github.com/suziepalmer10/MicrobiomeMetabolomeIntegration2026.

## Author’s Contributions

SNP, AYK, and XZ planned the research; SNP, XZ, JK, RW, and SG performed the experiments; SNP analyzed the data; AM and SG contributed technical support. SNP, AAM, CMZ, XZ, DL, and AYK wrote the paper.

## Acknowledgements

Not applicable

## Clinical Trial Number

Not applicable

## Supplemental Figures

**Supplemental Figure 1.**
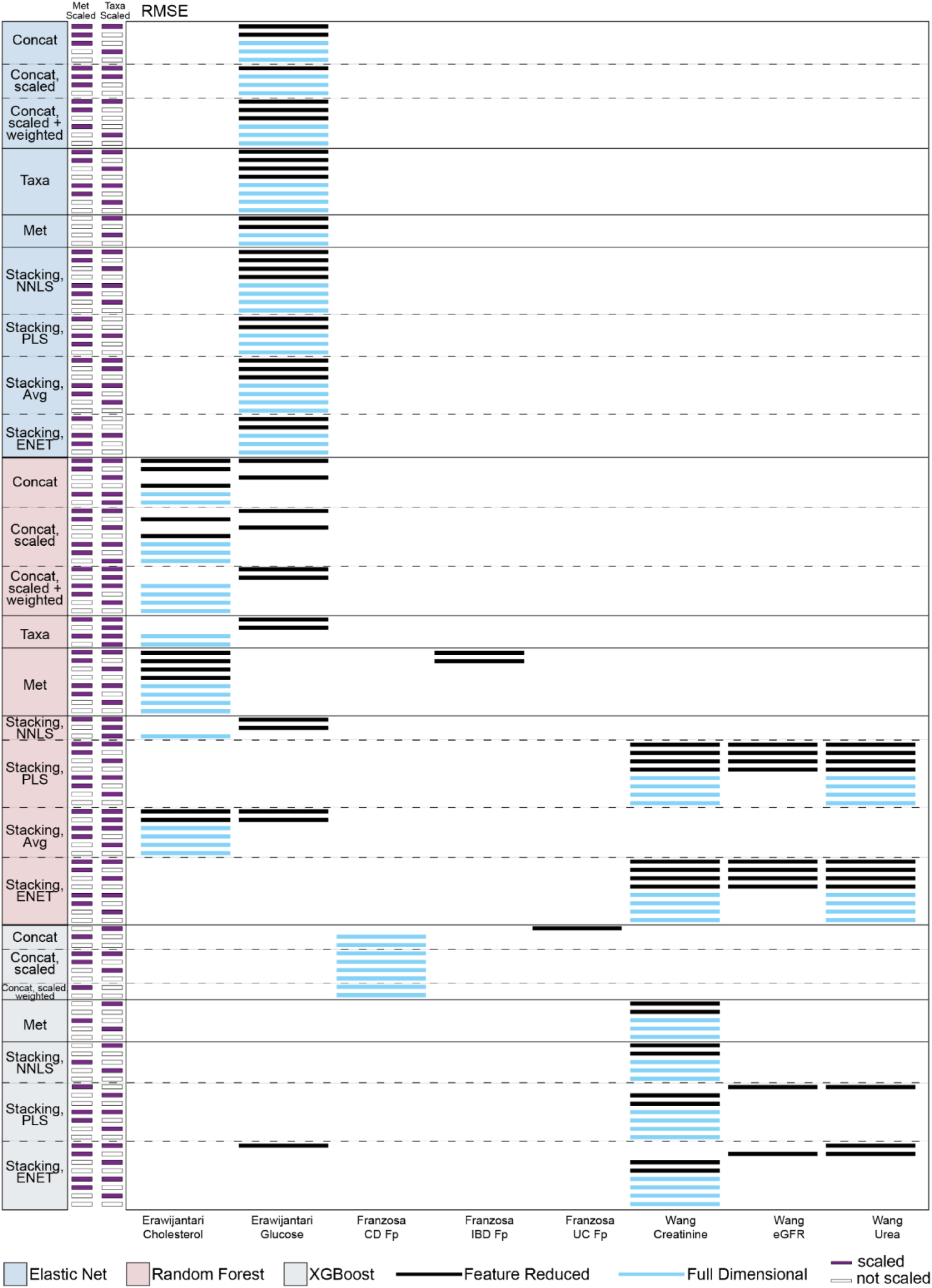
Dataset-level RMSE competitive sets support aggregated continuous outcome trends. Dataset-level competitive-set map for continuous prediction tasks evaluated by root mean squared error (RMSE). Columns represent individual continuous outcomes evaluated across the four datasets. Rows represent model configurations grouped by machine learning algorithm, omics input or integration strategy, feature scaling, and dimensionality setting. Competitive-set membership was defined within each endpoint using held-out test RMSE averaged across 20 repeated train/test splits, retaining configurations within 2% of the lowest mean RMSE. Side annotations indicate metabolomics and taxonomic scaling status (purple – scaled, white – unscaled), and horizontal marks indicate whether the retained configuration used feature-reduced (black) or full-dimensional (blue) predictors. ENET, Elastic Net; RF, Random Forest; XGB, XGBoost; Met, metabolomics; Taxa, taxonomic relative abundance; Concat, concatenation; CLR, centered log-ratio; NNLS, non-negative least squares; PLS, partial least squares; Avg, averaging; RMSE, root mean squared error.

**Supplemental Figure 2.**
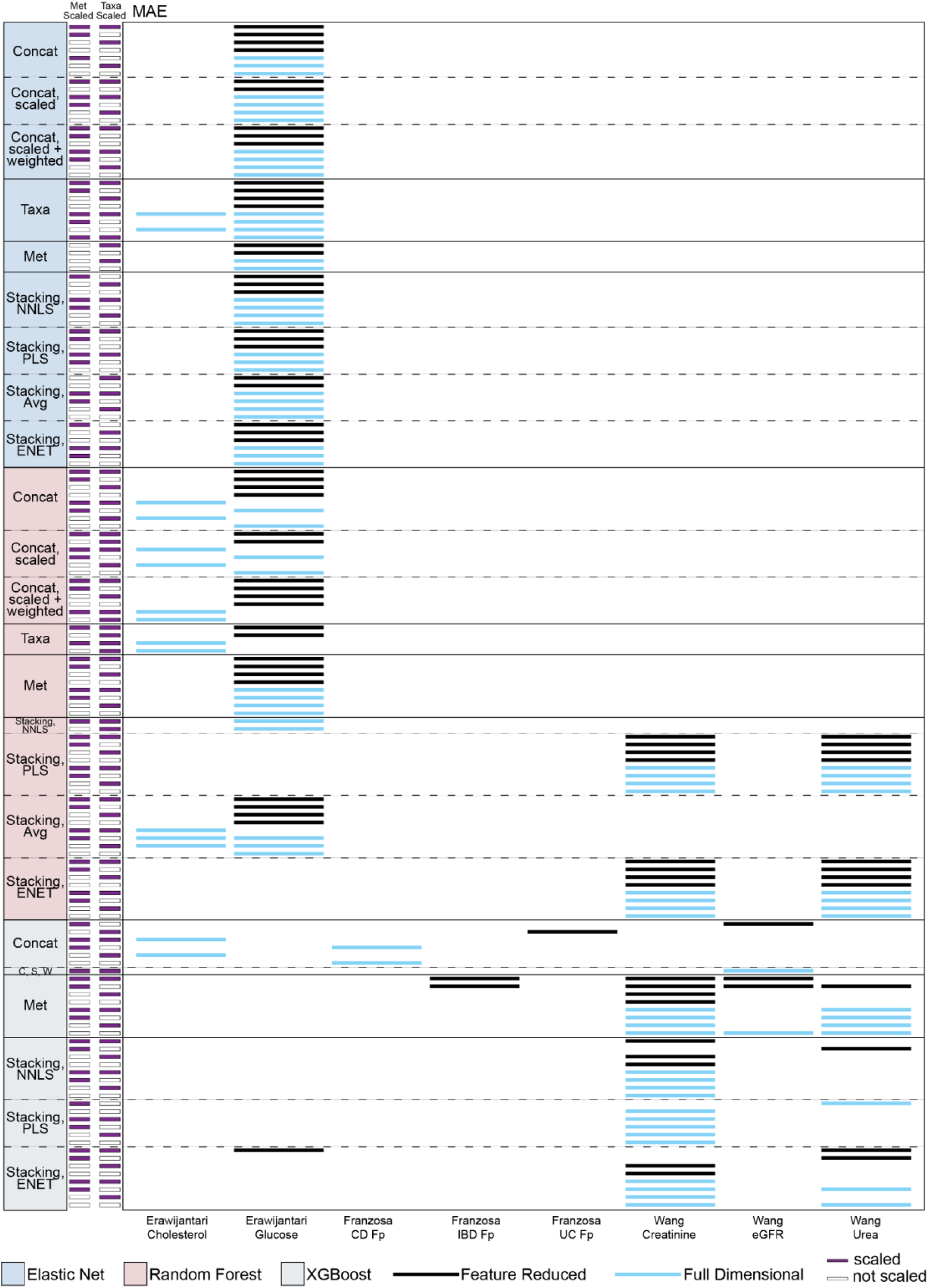
Dataset-level MAE competitive sets support aggregated continuous outcome trends. Dataset-level competitive-set map for continuous prediction tasks evaluated by mean absolute error (MAE). Columns represent the same continuous endpoints shown in Supplementary Figure 1, and rows represent model configurations grouped by machine learning algorithm, omics input or integration strategy, feature scaling, and dimensionality setting. Competitive-set membership was defined within each endpoint using held-out test MAE averaged across 20 repeated train/test splits, retaining configurations within 2% of the lowest mean MAE. Side annotations indicate metabolomics and taxonomic scaling status (purple – scaled, white – unscaled), and horizontal marks indicate whether the retained configuration used feature-reduced (black) or full-dimensional (blue) predictors. This map provides the endpoint-level competitive-set memberships underlying the aggregate MAE summaries in Figure 2. ENET, Elastic Net; RF, Random Forest; XGB, XGBoost; Met, metabolomics; Taxa, taxonomic relative abundance; Concat, concatenation; CLR, centered log-ratio; NNLS, non-negative least squares; PLS, partial least squares; Avg, averaging; MAE, mean absolute error.

**Supplemental Figure 3.**
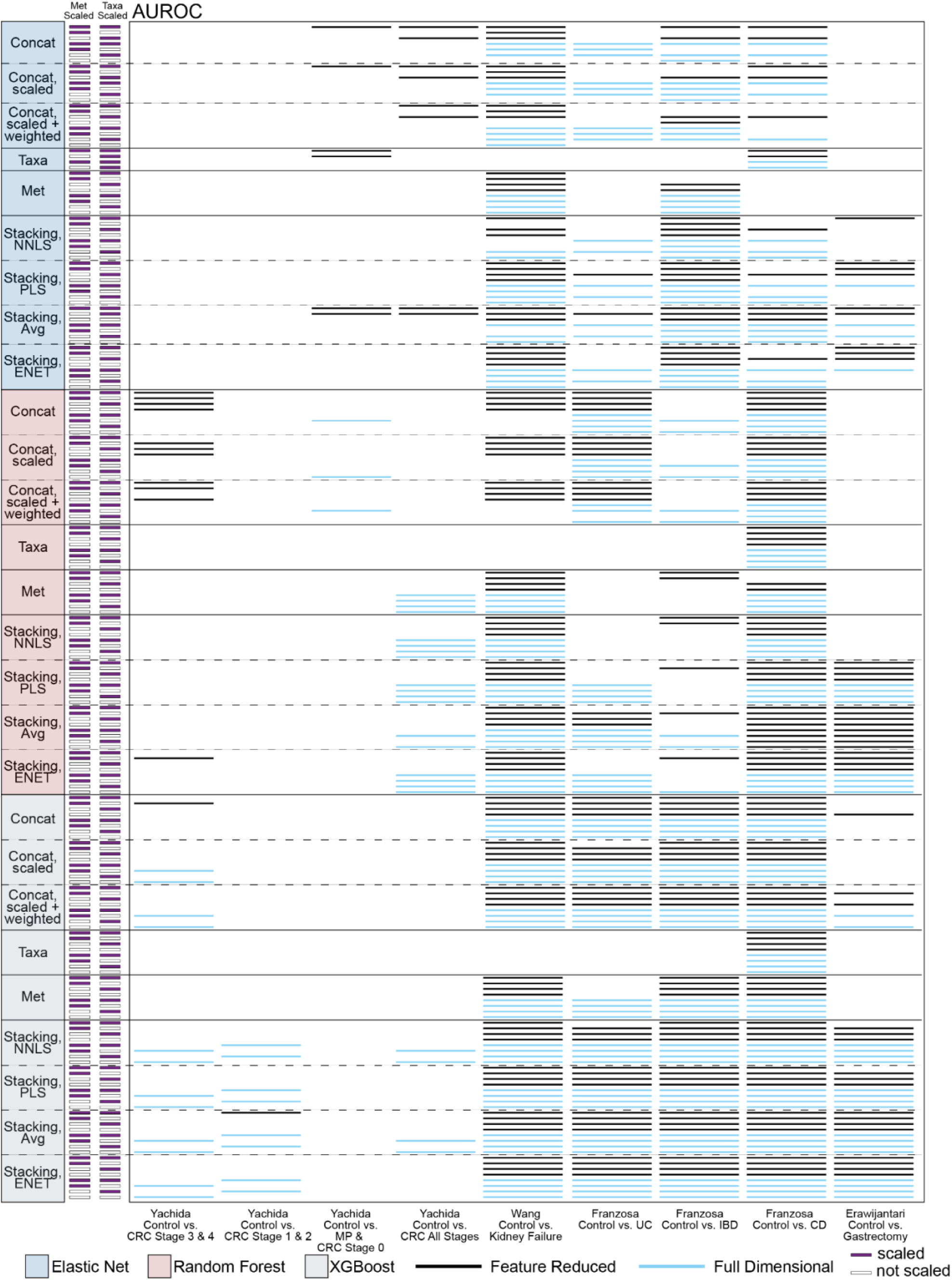
Dataset-level AUROC competitive sets support aggregated binary outcome trends. Dataset-level competitive-set map for binary classification tasks evaluated by area under the receiver operating characteristic curve (AUROC). Columns represent individual binary outcomes evaluated across the four datasets. Rows represent model configurations grouped by machine learning algorithm, omics input or integration strategy, feature scaling, and dimensionality setting. Competitive-set membership was defined within each endpoint using held-out test AUROC averaged across 20 repeated train/test splits, retaining configurations within 2% of the highest mean AUROC. Side annotations indicate metabolomics and taxonomic scaling status (purple – scaled, white – unscaled), and horizontal marks indicate whether the retained configuration used feature-reduced (black) or full-dimensional (blue) predictors. This map provides the endpoint-level competitive-set memberships underlying the aggregate AUROC summaries in Figure 3. ENET, Elastic Net; RF, Random Forest; XGB, XGBoost; Met, metabolomics; Taxa, taxonomic relative abundance; Concat, concatenation; CLR, centered log-ratio; NNLS, non-negative least squares; PLS, partial least squares; Avg, averaging; AUROC, area under the receiver operating characteristic curve; UC, ulcerative colitis; IBD, inflammatory bowel disease; CD, Crohn’s disease; CRC, colorectal cancer; MP, multiple polypoid adenomas.

**Supplemental Figure 4.**
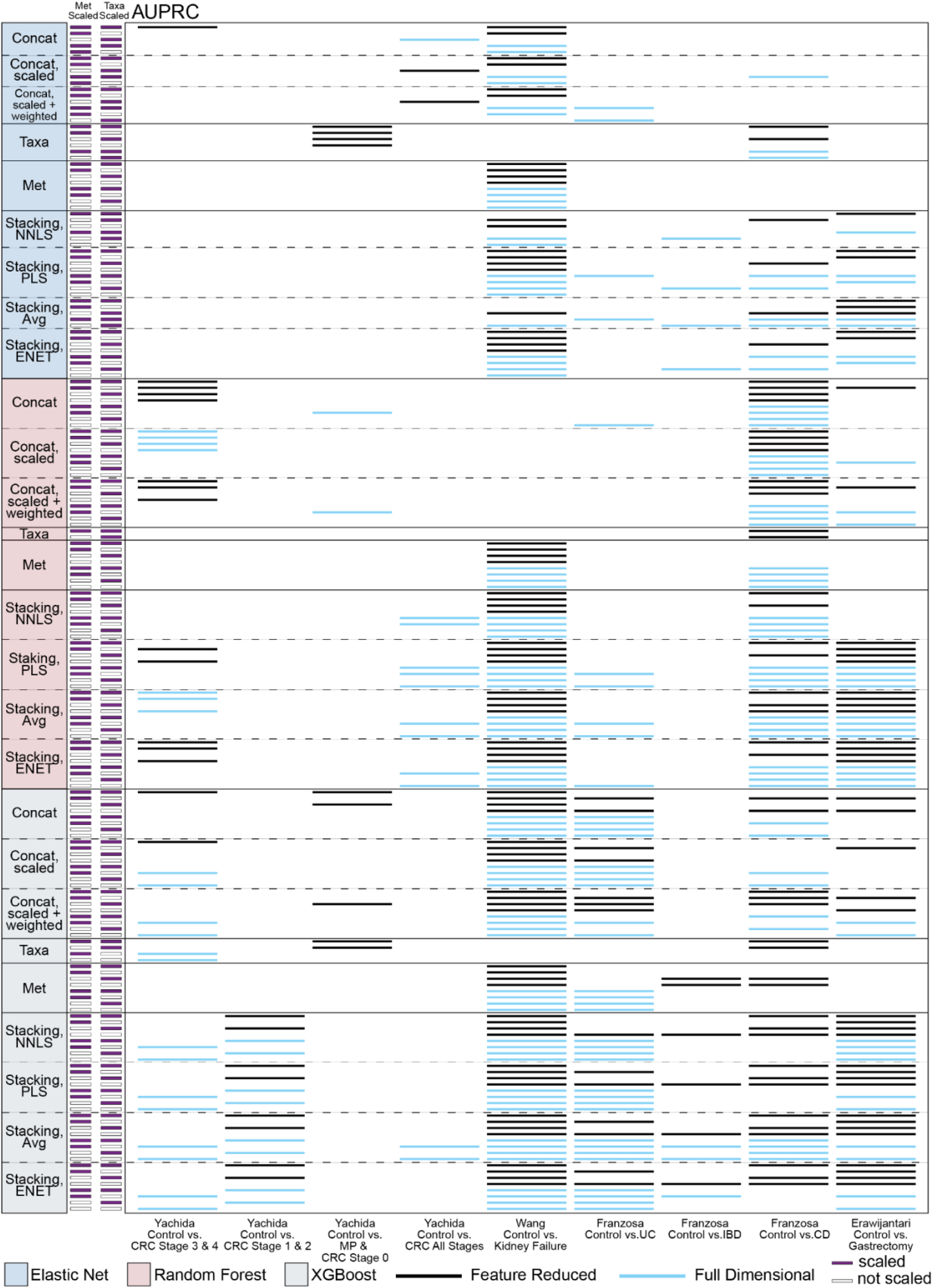
Dataset-level AUPRC competitive sets support aggregated binary outcome trends. Dataset-level competitive-set map for binary classification tasks evaluated by area under the precision-recall curve (AUPRC). Columns represent the same binary endpoints shown in Supplementary Figure 3, and rows represent model configurations grouped by machine learning algorithm, omics input or integration strategy, feature scaling, and dimensionality setting. Competitive-set membership was defined within each endpoint using held-out test AUPRC averaged across 20 repeated train/test splits, retaining configurations within 2% of the highest mean AUPRC. Side annotations indicate metabolomics and taxonomic scaling status (purple – scaled, white – unscaled), and horizontal marks indicate whether the retained configuration used feature-reduced (black) or full-dimensional (blue) predictors. This map provides the endpoint-level competitive-set memberships underlying the aggregate AUPRC summaries in Figure 3. ENET, Elastic Net; RF, Random Forest; XGB, XGBoost; Met, metabolomics; Taxa, taxonomic relative abundance; Concat, concatenation; CLR, centered log-ratio; NNLS, non-negative least squares; PLS, partial least squares; Avg, averaging; AUPRC, area under the precision-recall curve; UC, ulcerative colitis; IBD, inflammatory bowel disease; CD, Crohn’s disease; CRC, colorectal cancer; MP, multiple polypoid adenomas.

**Supplemental Figure 5.**
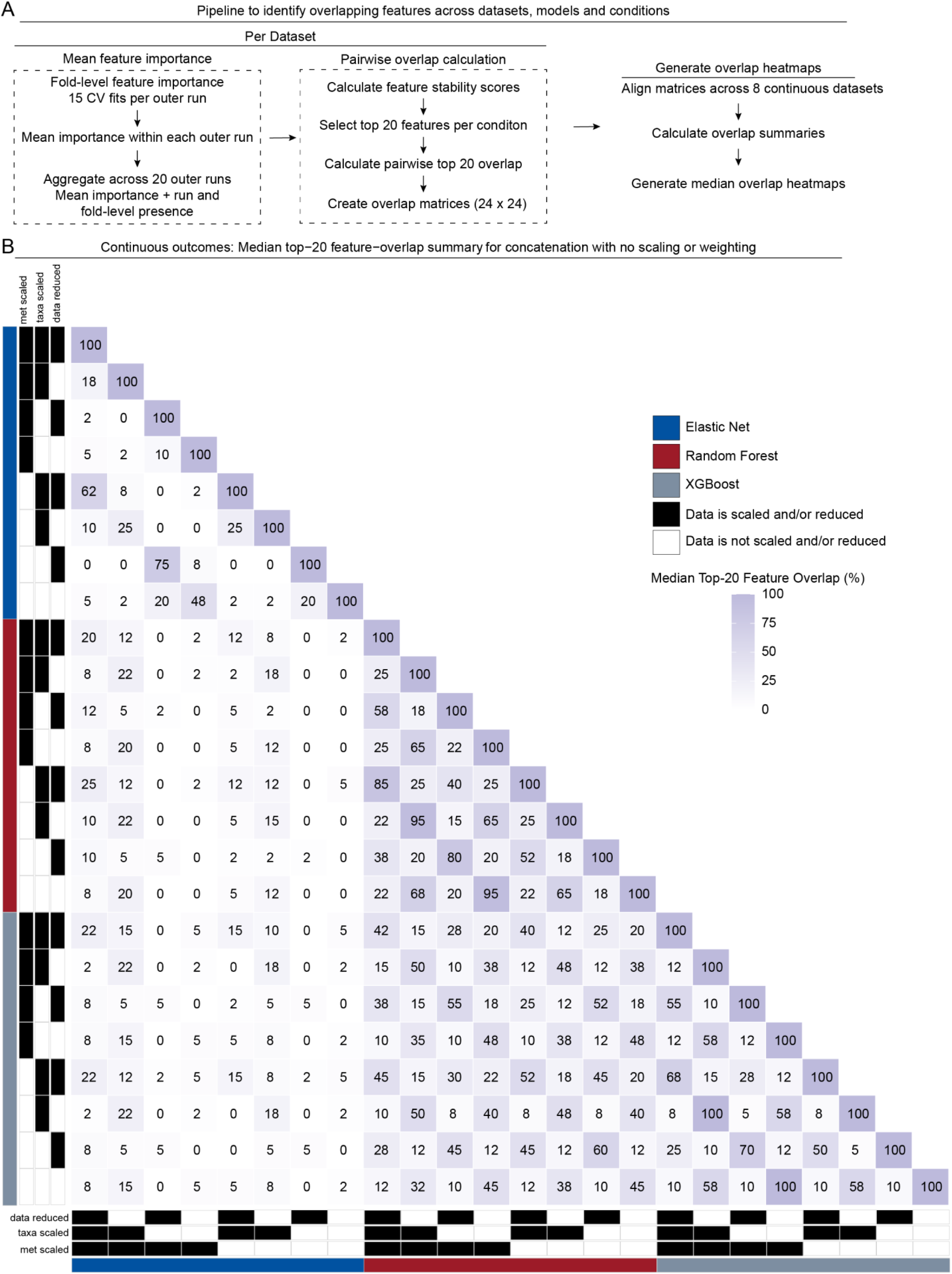
Median top 20 feature overlap across modeling and preprocessing conditions in concatenated data for continuous outcomes. **(A)** Workflow used to quantify feature-selection stability and overlap. For each study–outcome dataset, data view, and pipeline condition, feature importance was extracted from the 15 cross-validation fits within each of 20 outer runs, corresponding to three repeats of five-fold cross-validation per run. Fold-level importance values were averaged within each outer run and then across the outer runs in which the feature was present. Features absent from a fold or run were not assigned an importance of zero but contributed to the corresponding presence-frequency. Mean importance was converted to a percentile rank, and the primary feature stability score was calculated as the importance percentile rank multiplied by outer-run presence frequency. A secondary score based on fold level presence frequency contributed to the prespecified tie-breaking procedure. Features were ranked by the primary stability score, and the 20 highest-ranking features were retained for each condition Pairwise overlap was calculated (See Methods) producing a 24 × 24 overlap matrix for each concatenated view. Corresponding condition pairs were aligned across 8 datasets with continuous outcomes, and overlap values were summarized across datasets. **(B)** Median top 20 feature-overlap matrices for the concatenated view, with no scaling or weighting, for continuous outcomes. Each row and column represents one of 24 pipeline conditions defined by three prediction models (elastic net, random forest, and XGBoost) and eight view-specific preprocessing combinations.

**Supplemental Figure 6.**
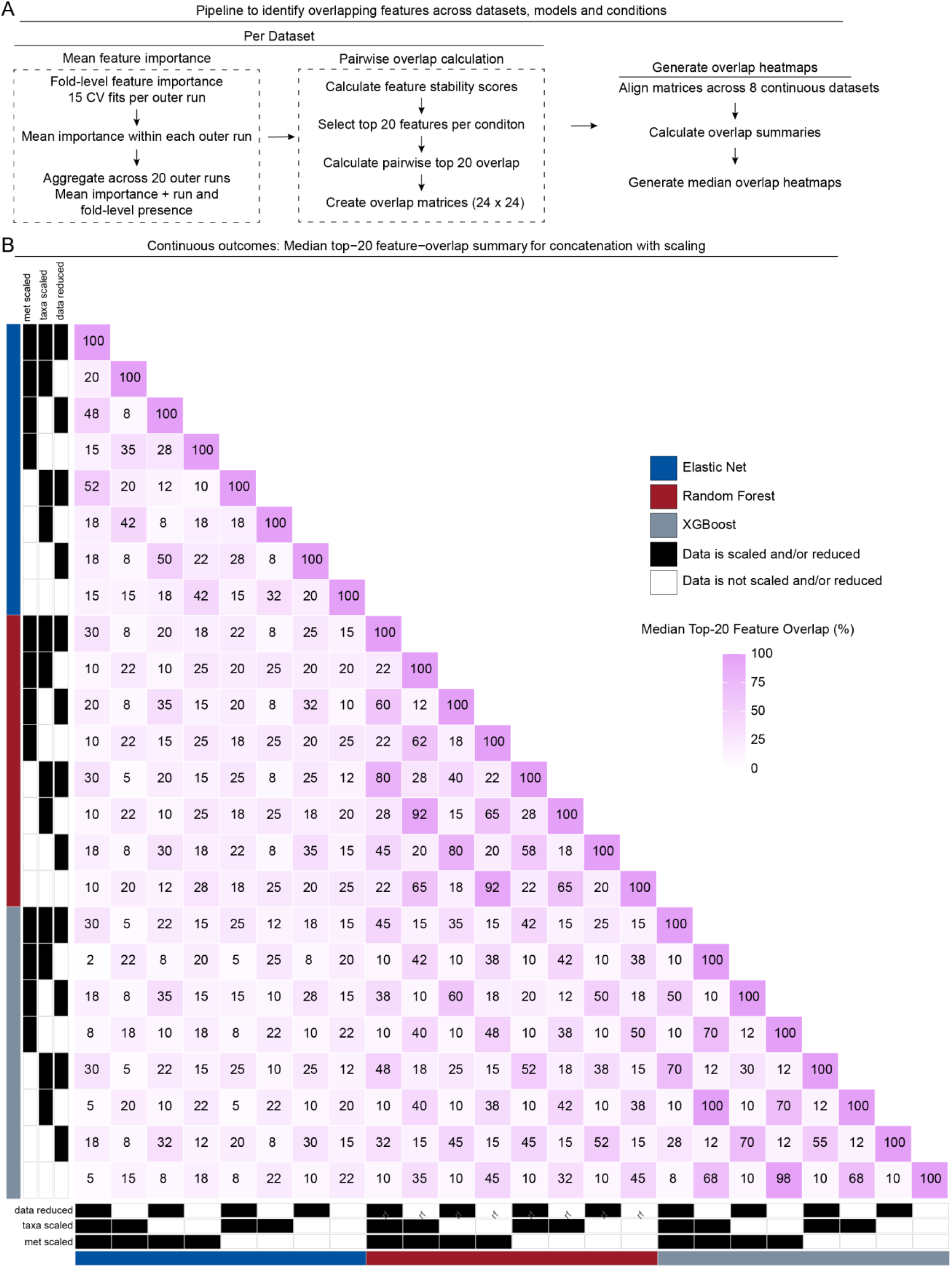
Median top 20 feature overlap across modeling and preprocessing conditions in z-score scaled concatenated data for continuous outcomes. **(A)** Workflow used to quantify feature-selection stability and overlap. For each study–outcome dataset, data view, and pipeline condition, feature importance was extracted from the 15 cross-validation fits within each of 20 outer runs, corresponding to three repeats of five-fold cross-validation per run. Fold-level importance values were averaged within each outer run and then across the outer runs in which the feature was present. Features absent from a fold or run were not assigned an importance of zero but contributed to the corresponding presence-frequency. Mean importance was converted to a percentile rank, and the primary feature stability score was calculated as the importance percentile rank multiplied by outer-run presence frequency. A secondary score based on fold level presence frequency contributed to the prespecified tie-breaking procedure. Features were ranked by the primary stability score, and the 20 highest-ranking features were retained for each condition Pairwise overlap was calculated (See Methods) producing a 24 × 24 overlap matrix for each concatenated view. Corresponding condition pairs aligned across 8 datasets with continuous outcomes, and overlap values were summarized across datasets. **(B)** Median top 20 feature-overlap matrices for the concatenated view, with z-score scaling for continuous outcomes. Each row and column represents one of 24 pipeline conditions defined by three prediction models (elastic net, random forest, and XGBoost) and eight view-specific preprocessing combinations.

**Supplemental Figure 7.**
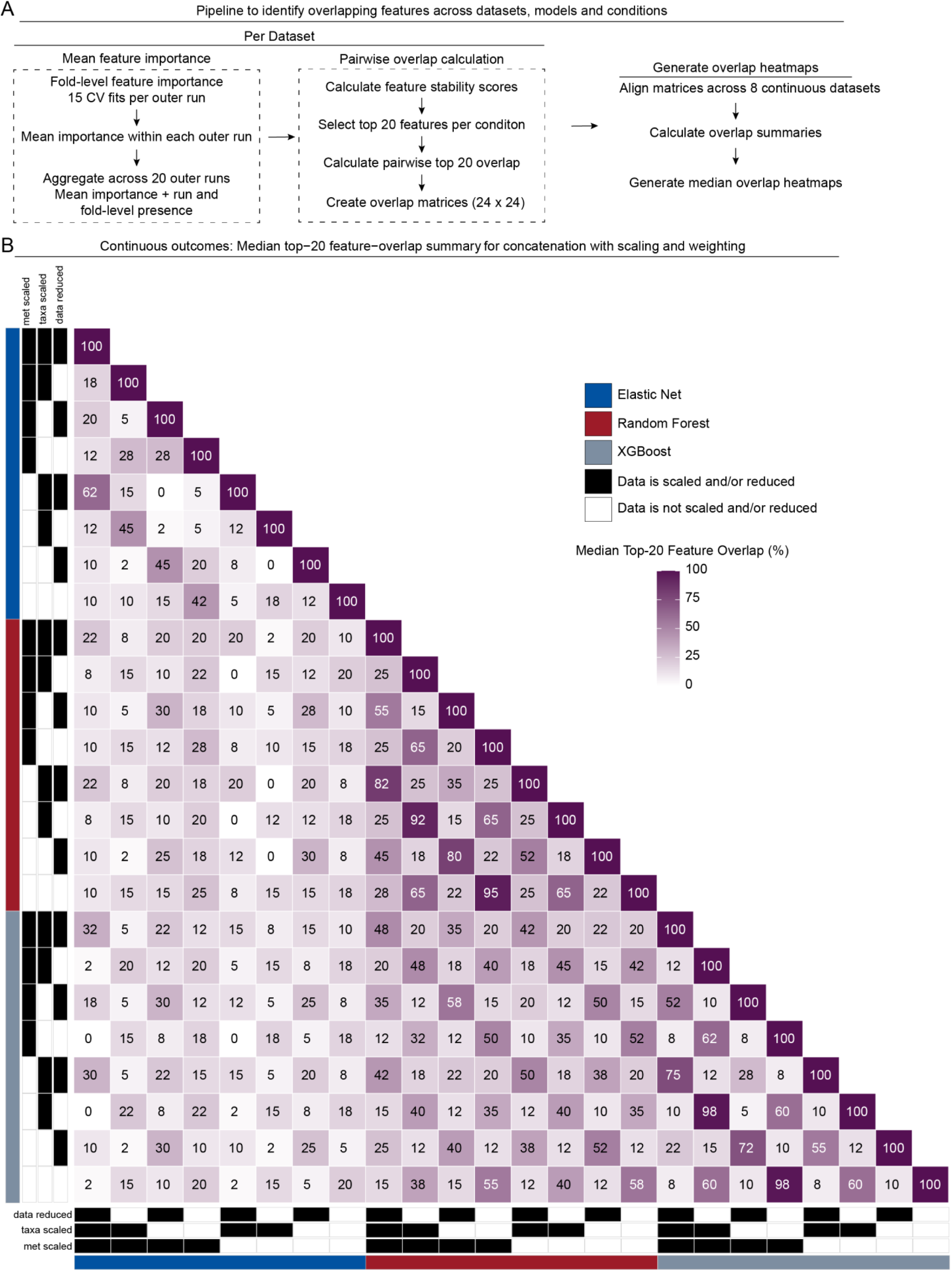
Median top 20 feature overlap across modeling and preprocessing conditions in z-score scaled, block weighted concatenated data for continuous outcomes. **(A)** Workflow used to quantify feature-selection stability and overlap. For each study–outcome dataset, data view, and pipeline condition, feature importance was extracted from the 15 cross-validation fits within each of 20 outer runs, corresponding to three repeats of five-fold cross-validation per run. Fold-level importance values were averaged within each outer run and then across the outer runs in which the feature was present. Features absent from a fold or run were not assigned an importance of zero but contributed to the corresponding presence-frequency. Mean importance was converted to a percentile rank, and the primary feature stability score was calculated as the importance percentile rank multiplied by outer-run presence frequency. A secondary score based on fold level presence frequency contributed to the prespecified tie-breaking procedure. Features were ranked by the primary stability score, and the 20 highest-ranking features were retained for each condition Pairwise overlap was calculated (See Methods) producing a 24 × 24 overlap matrix for each concatenated view. Corresponding condition pairs aligned across 8 datasets with continuous outcomes, and overlap values were summarized across datasets. **(B)** Median top 20 feature-overlap matrices for the concatenated view, with z-score scaling and block weighting, for continuous outcomes. Each row and column represents one of 24 pipeline conditions defined by three prediction models (elastic net, random forest, and XGBoost) and eight view-specific preprocessing combinations.

**Supplemental Figure 8.**
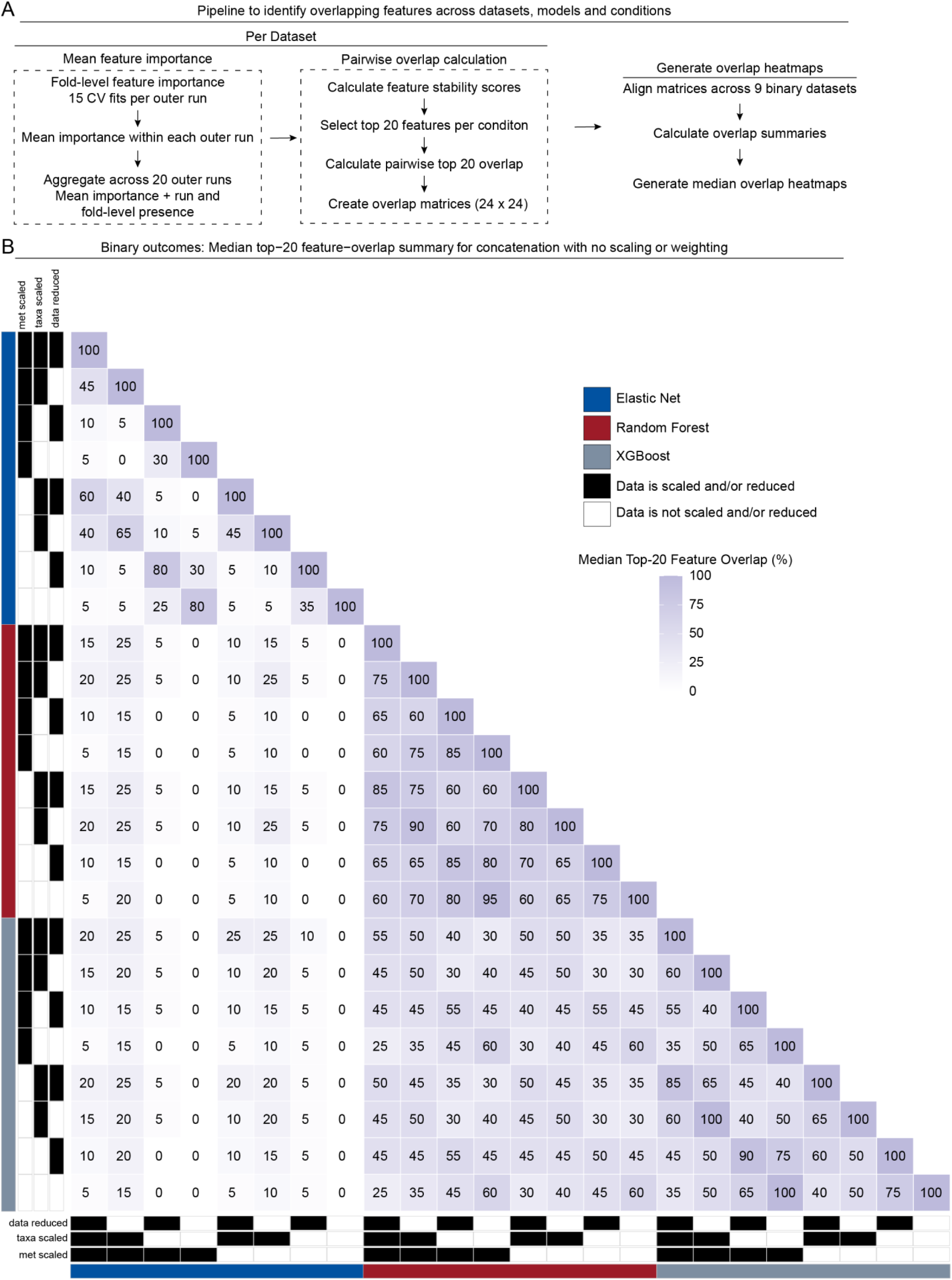
Median top 20 feature overlap across modeling and preprocessing conditions in concatenated data for binary outcomes **(A)** Workflow used to quantify feature-selection stability and overlap. For each study–outcome dataset, data view, and pipeline condition, feature importance was extracted from the 15 cross-validation fits within each of 20 outer runs, corresponding to three repeats of five-fold cross-validation per run. Fold-level importance values were averaged within each outer run and then across the outer runs in which the feature was present. Features absent from a fold or run were not assigned an importance of zero but contributed to the corresponding presence-frequency. Mean importance was converted to a percentile rank, and the primary feature stability score was calculated as the importance percentile rank multiplied by outer-run presence frequency. A secondary score based on fold level presence frequency contributed to the prespecified tie-breaking procedure. Features were ranked by the primary stability score, and the 20 highest-ranking features were retained for each condition Pairwise overlap was calculated (See Methods) producing a 24 × 24 overlap matrix for each concatenated view. Corresponding condition pairs were aligned across 9 datasets with binary outcomes, and overlap values were summarized across datasets. **(B)** Median top 20 feature-overlap matrices for the concatenated view, with no scaling or weighting, for continuous outcomes. Each row and column represents one of 24 pipeline conditions defined by three prediction models (elastic net, random forest, and XGBoost) and eight view-specific preprocessing combinations.

**Supplemental Figure 9.**
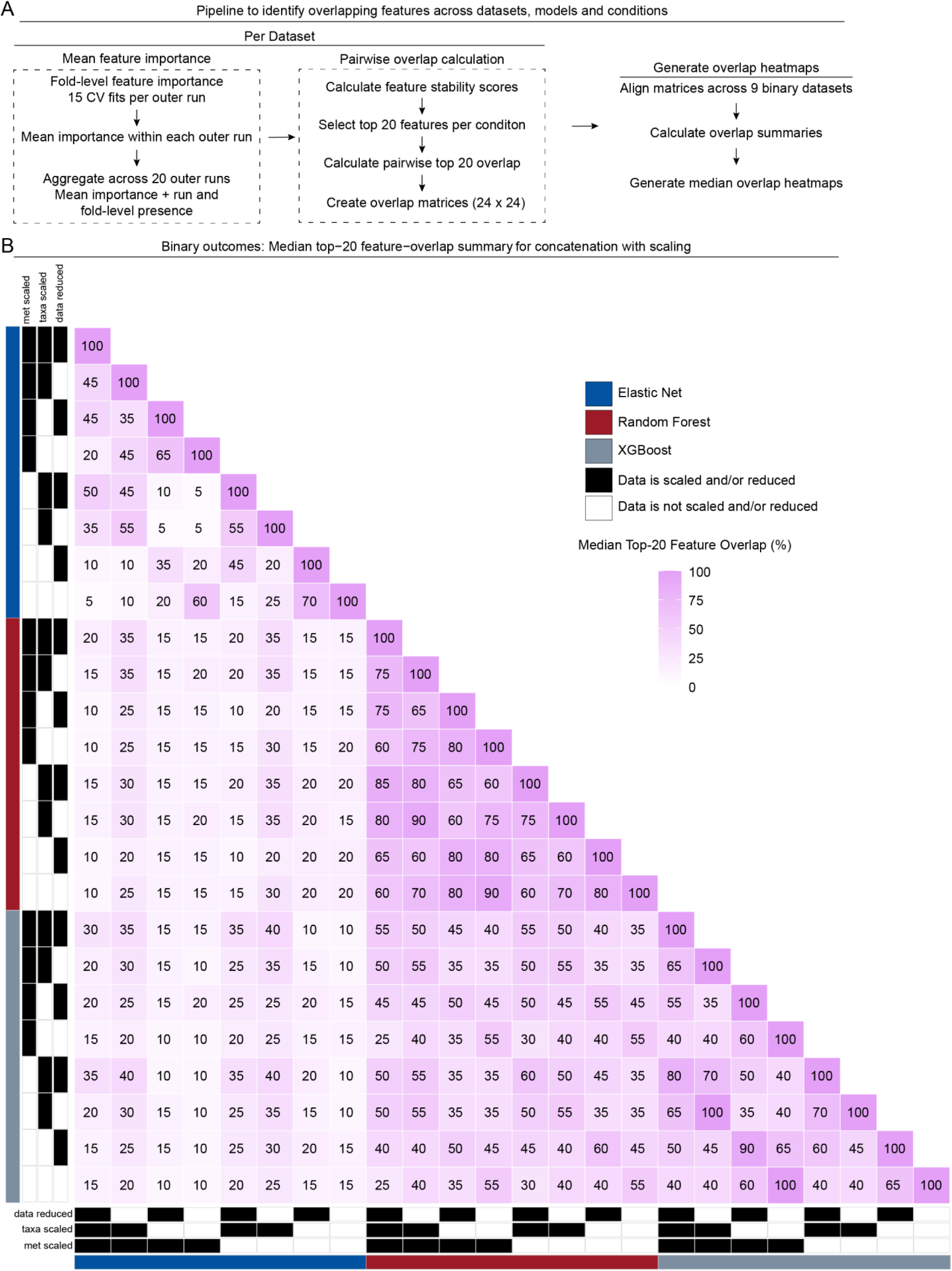
Median top 20 feature overlap across modeling and preprocessing conditions in z-score scaled concatenated data for binary outcomes. **(A)** Workflow used to quantify feature-selection stability and overlap. For each study–outcome dataset, data view, and pipeline condition, feature importance was extracted from the 15 cross-validation fits within each of 20 outer runs, corresponding to three repeats of five-fold cross-validation per run. Fold-level importance values were averaged within each outer run and then across the outer runs in which the feature was present. Features absent from a fold or run were not assigned an importance of zero but contributed to the corresponding presence-frequency. Mean importance was converted to a percentile rank, and the primary feature stability score was calculated as the importance percentile rank multiplied by outer-run presence frequency. A secondary score based on fold level presence frequency contributed to the prespecified tie-breaking procedure. Features were ranked by the primary stability score, and the 20 highest-ranking features were retained for each condition Pairwise overlap was calculated (See Methods) producing a 24 × 24 overlap matrix for each concatenated view. Corresponding condition pairs were aligned across 9 datasets with binary outcomes, and overlap values were summarized across datasets. **(B)** Median top 20 feature-overlap matrices for the concatenated view, with z-score scaling for continuous outcomes. Each row and column represents one of 24 pipeline conditions defined by three prediction models (elastic net, random forest, and XGBoost) and eight view-specific preprocessing combinations.

**Supplemental Figure 10.**
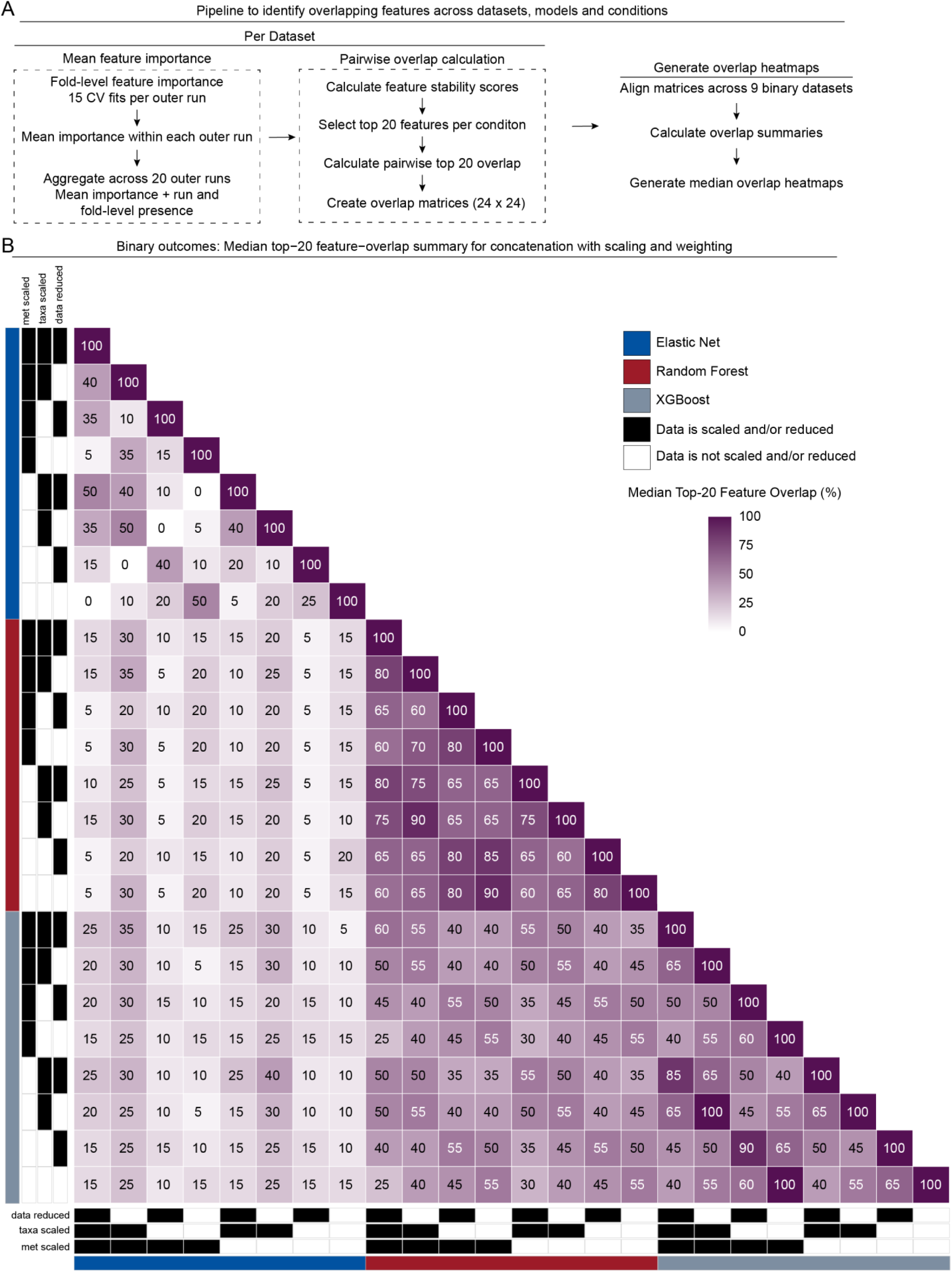
Median top 20 feature overlap across modeling and preprocessing conditions in z-score scaled, block weighted concatenated data for binary outcomes. **(A)** Workflow used to quantify feature-selection stability and overlap. For each study–outcome dataset, data view, and pipeline condition, feature importance was extracted from the 15 cross-validation fits within each of 20 outer runs, corresponding to three repeats of five-fold cross-validation per run. Fold-level importance values were averaged within each outer run and then across the outer runs in which the feature was present. Features absent from a fold or run were not assigned an importance of zero but contributed to the corresponding presence-frequency. Mean importance was converted to a percentile rank, and the primary feature stability score was calculated as the importance percentile rank multiplied by outer-run presence frequency. A secondary score based on fold level presence frequency contributed to the prespecified tie-breaking procedure. Features were ranked by the primary stability score, and the 20 highest-ranking features were retained for each condition Pairwise overlap was calculated (See Methods) producing a 24 × 24 overlap matrix for each concatenated view. Corresponding condition pairs were aligned across 9 datasets with binary outcomes, and overlap values were summarized across datasets. **(B)** Median top 20 feature-overlap matrices for the concatenated view, with z-score scaling and block weighting, for continuous outcomes. Each row and column represents one of 24 pipeline conditions defined by three prediction models (elastic net, random forest, and XGBoost) and eight view-specific preprocessing combinations.

