## Additional File 1 for "Trends in Machine Learning and Feature Selection Stability for Human Gut Microbiome (Shotgun Metagenomics) and Metabolomics Matched Datasets"

### Erawijantari Control vs. Gastrectomy Mean Performance with 95% CI Across Seeds
(● Data is scaled and/or reduced)

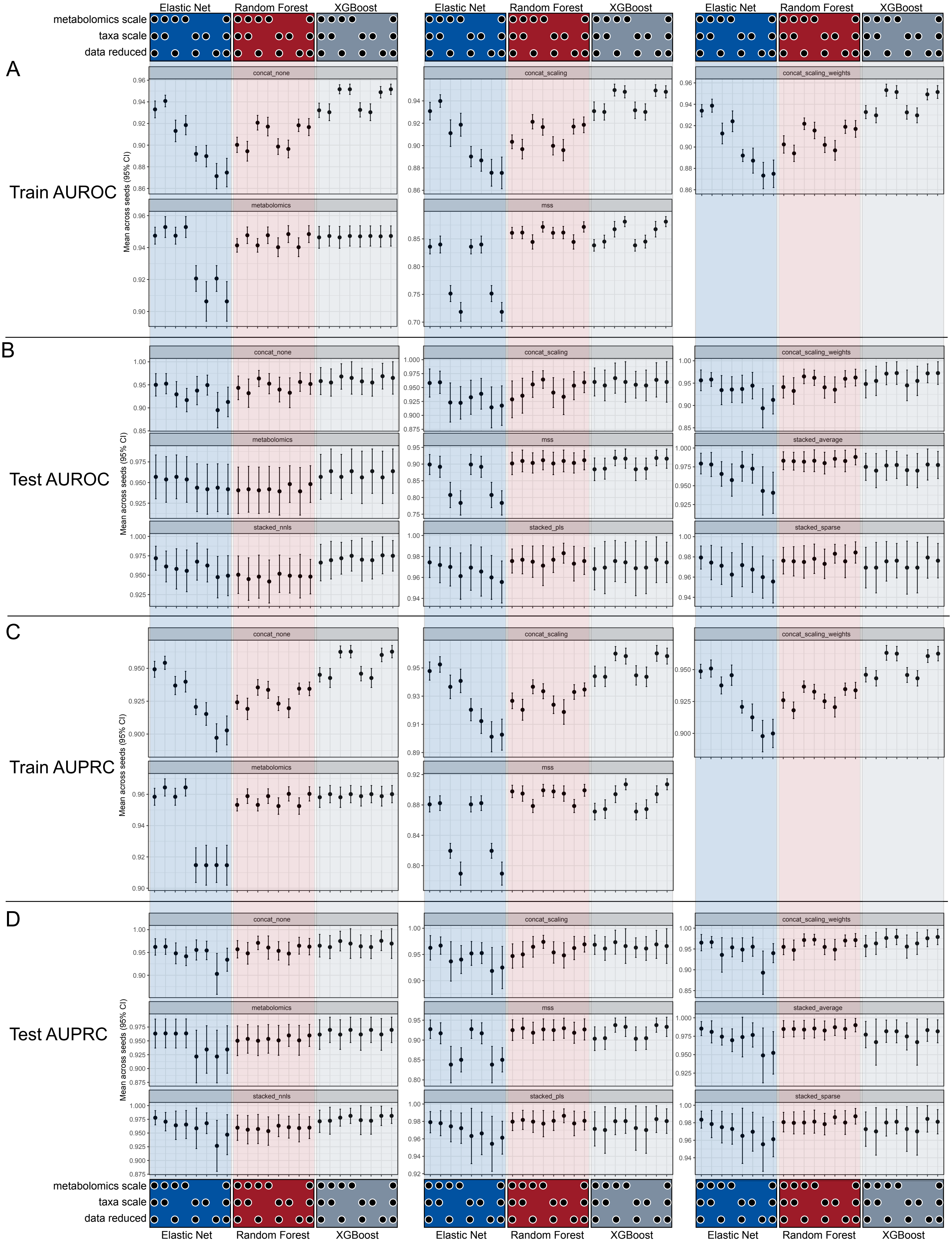

Erawijantari Cholesterol Mean Performance with 95% CI Across Seeds (● Data is scaled and/or reduced)

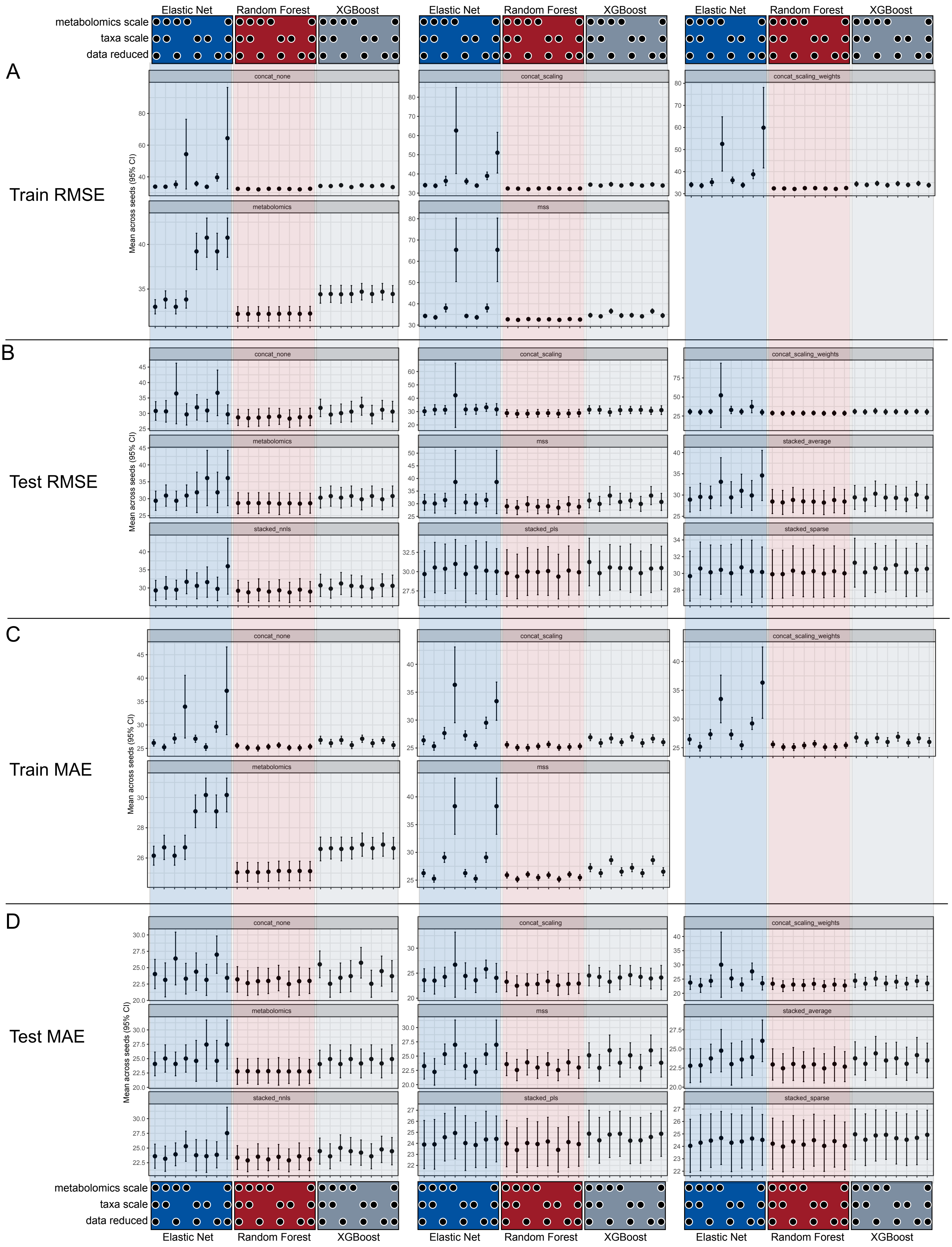

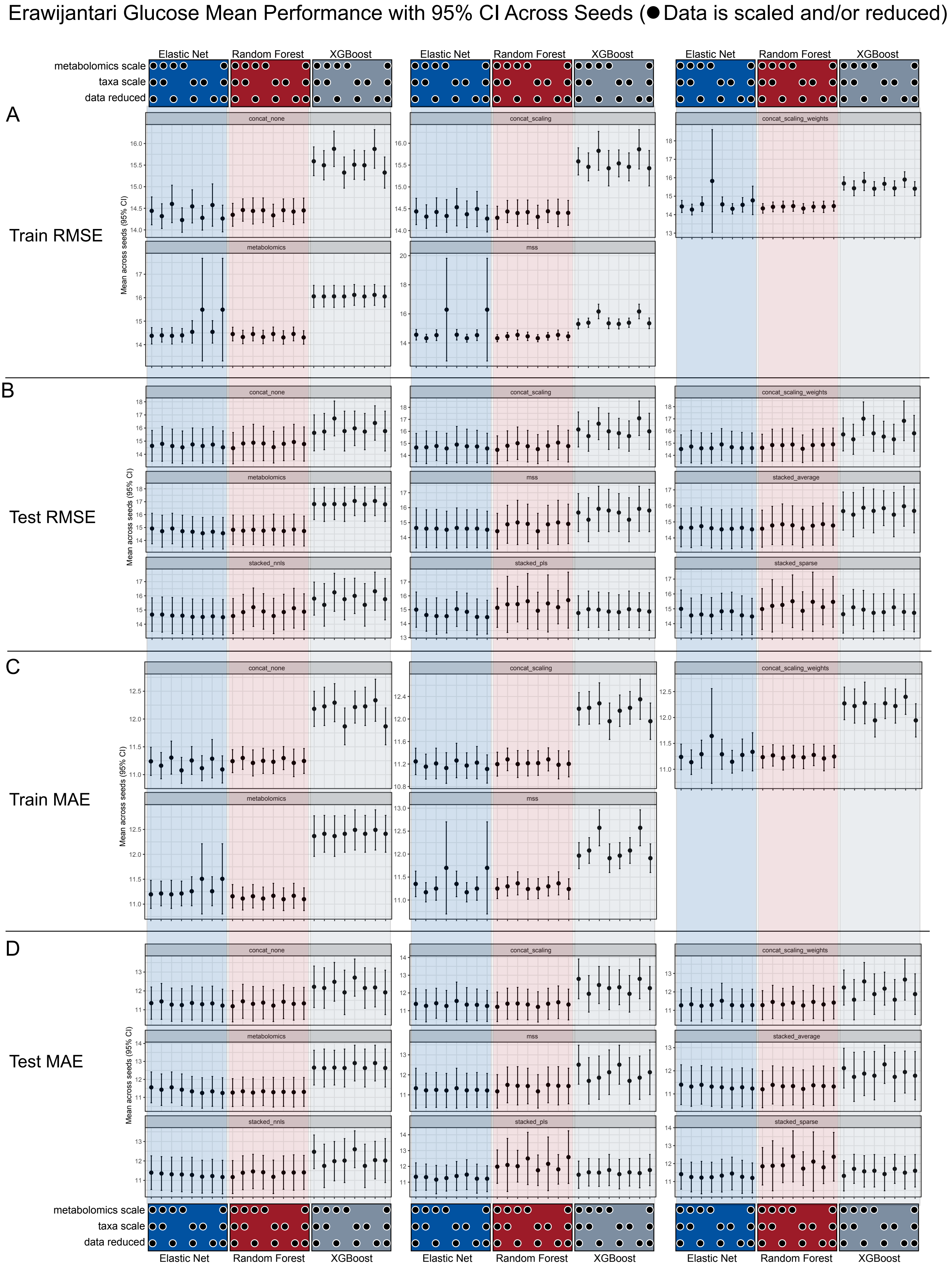

Erawijantari Glucose Mean Performance with 95% CI Across Seeds (● Data is scaled and/or reduced)
Elastic Net
Random Forest
XGBoost
metabolomics scale
taxa scale
data reduced
A
Train RMSE
B
Test RMSE
C
Train MAE
D
Test MAE
Mean across seeds (95% CI)
concat_none
concat_scaling
concat_scaling_weights
metabolomics
mss
stacked_average
stacked_nnls
stacked_pls
stacked_sparse

Franzosa Control vs. CD Mean Performance with 95% CI Across Seeds
(● Data is scaled and/or reduced)

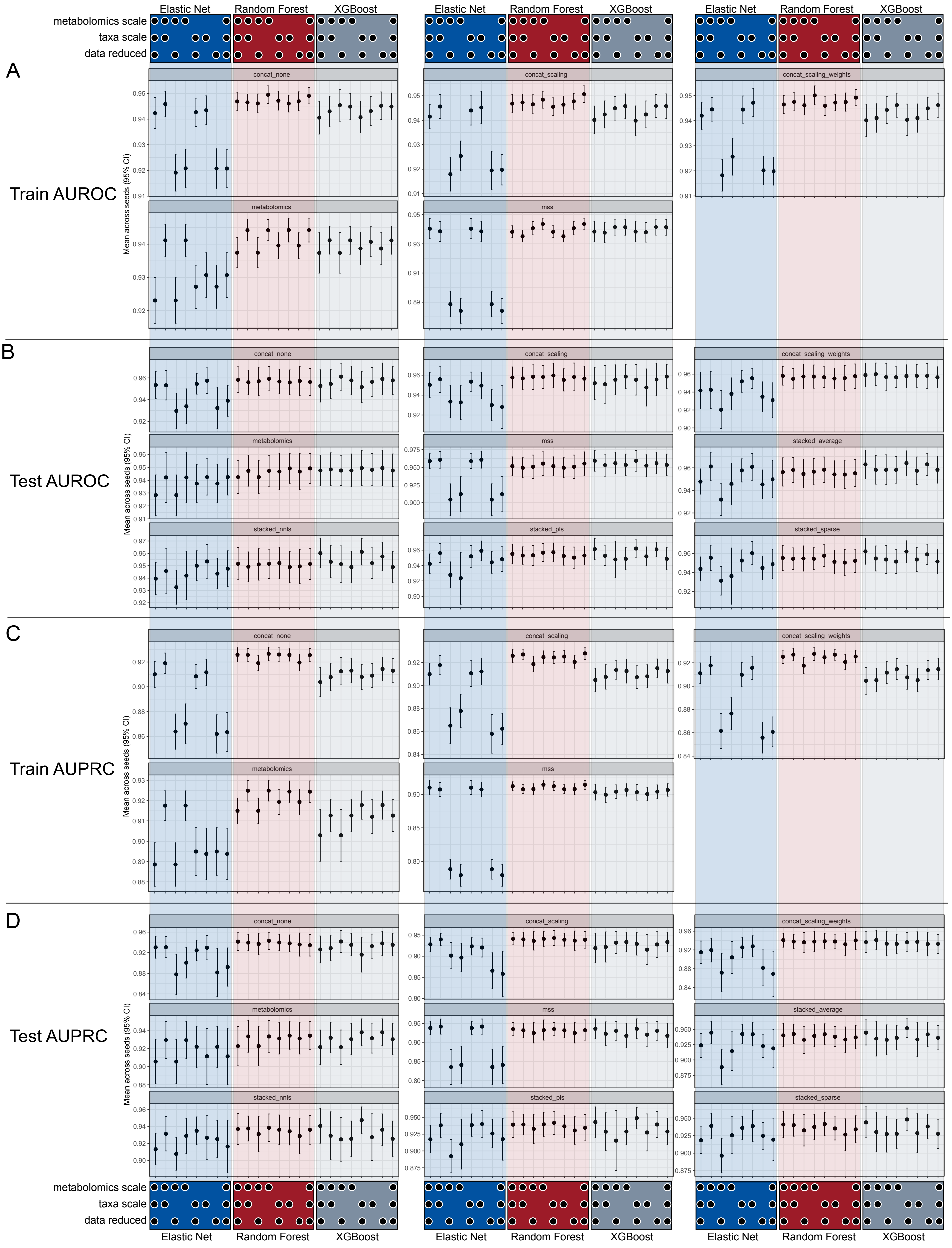

Franzosa CD Fp Mean Performance with 95% CI Across Seeds (● Data is scaled and/or reduced)

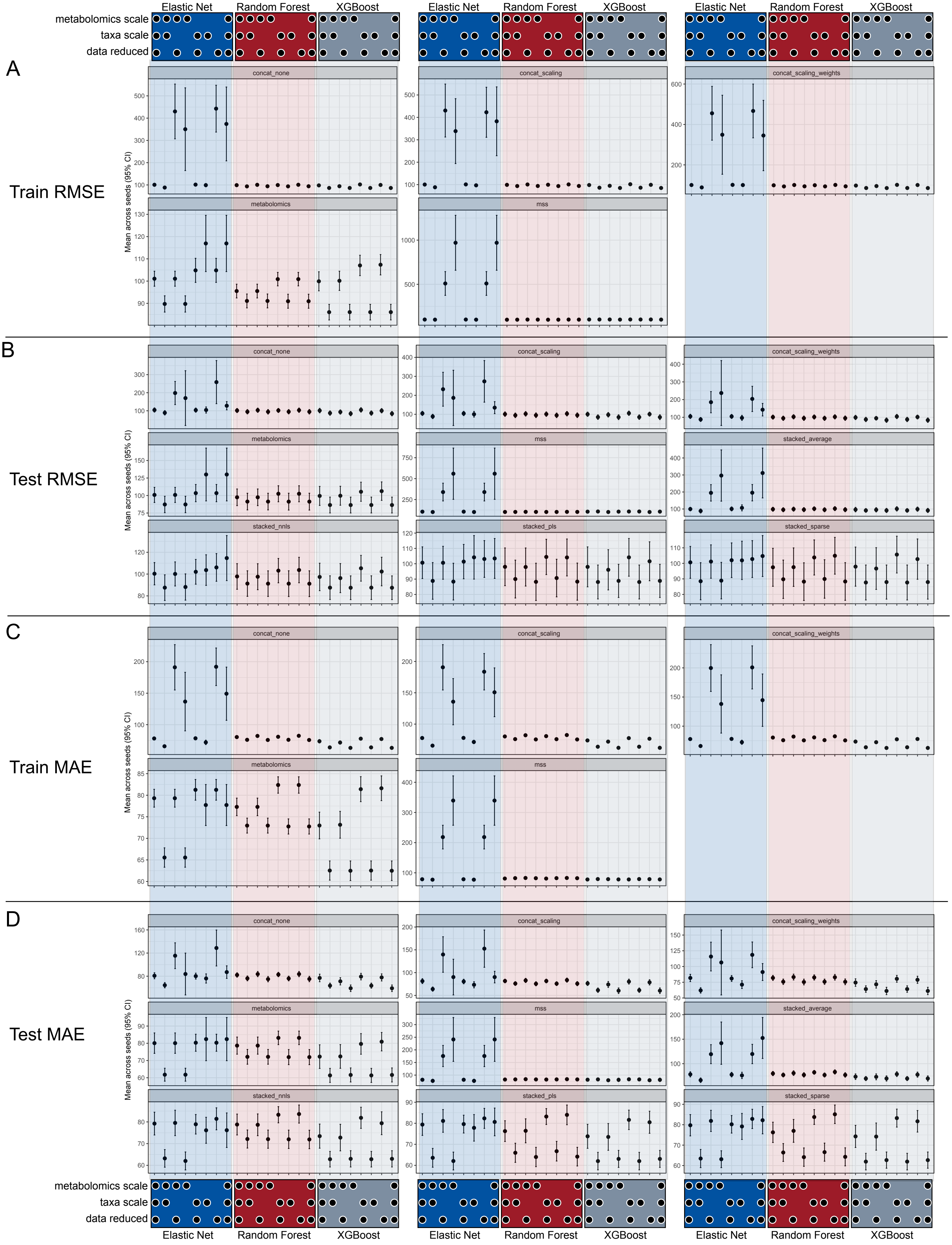

Franzosa Control vs. IBD Mean Performance with 95% CI Across Seeds
(● Data is scaled and/or reduced)

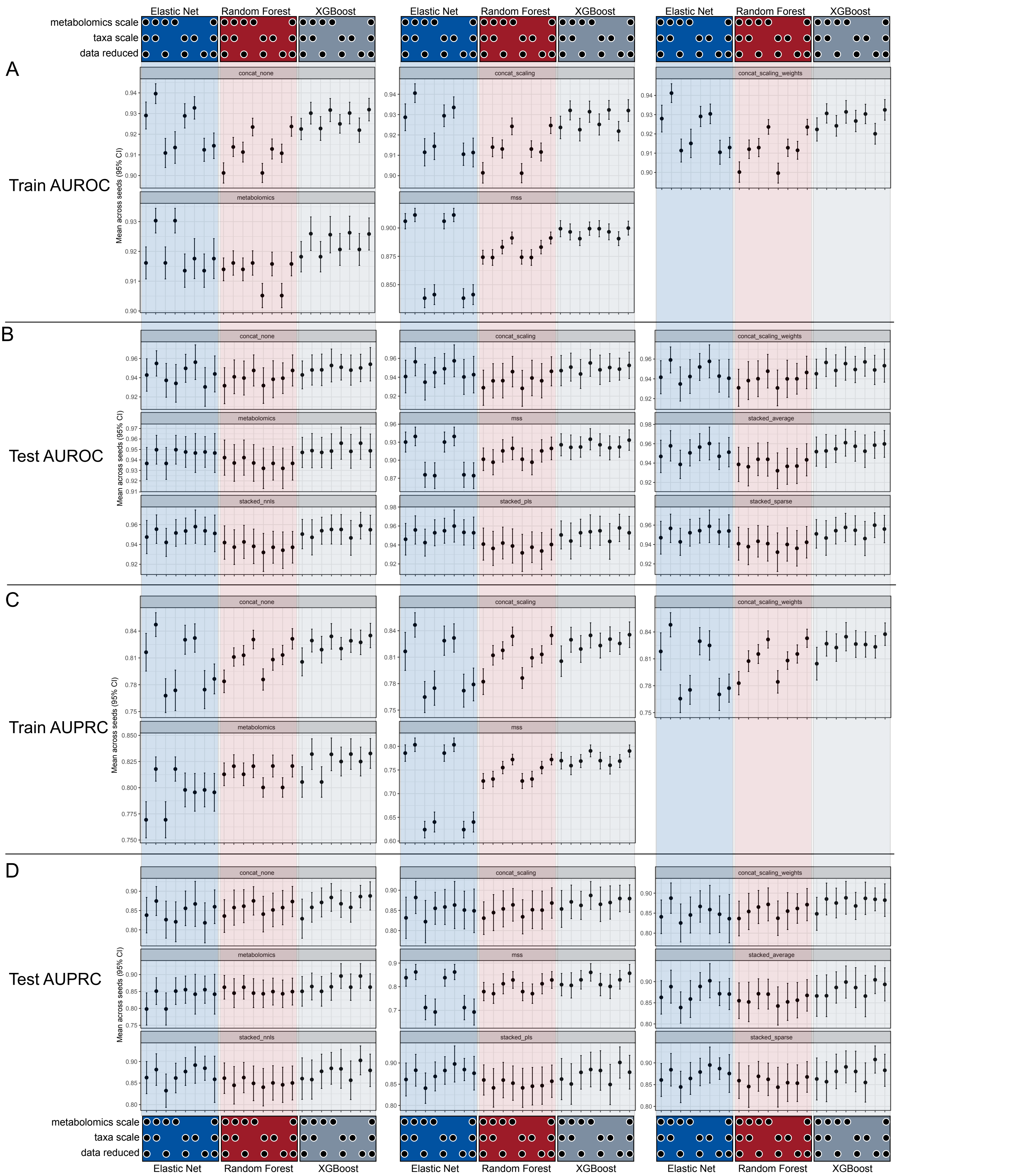

Franzosa IBD Fp Mean Performance with 95% CI Across Seeds (● Data is scaled and/or reduced)

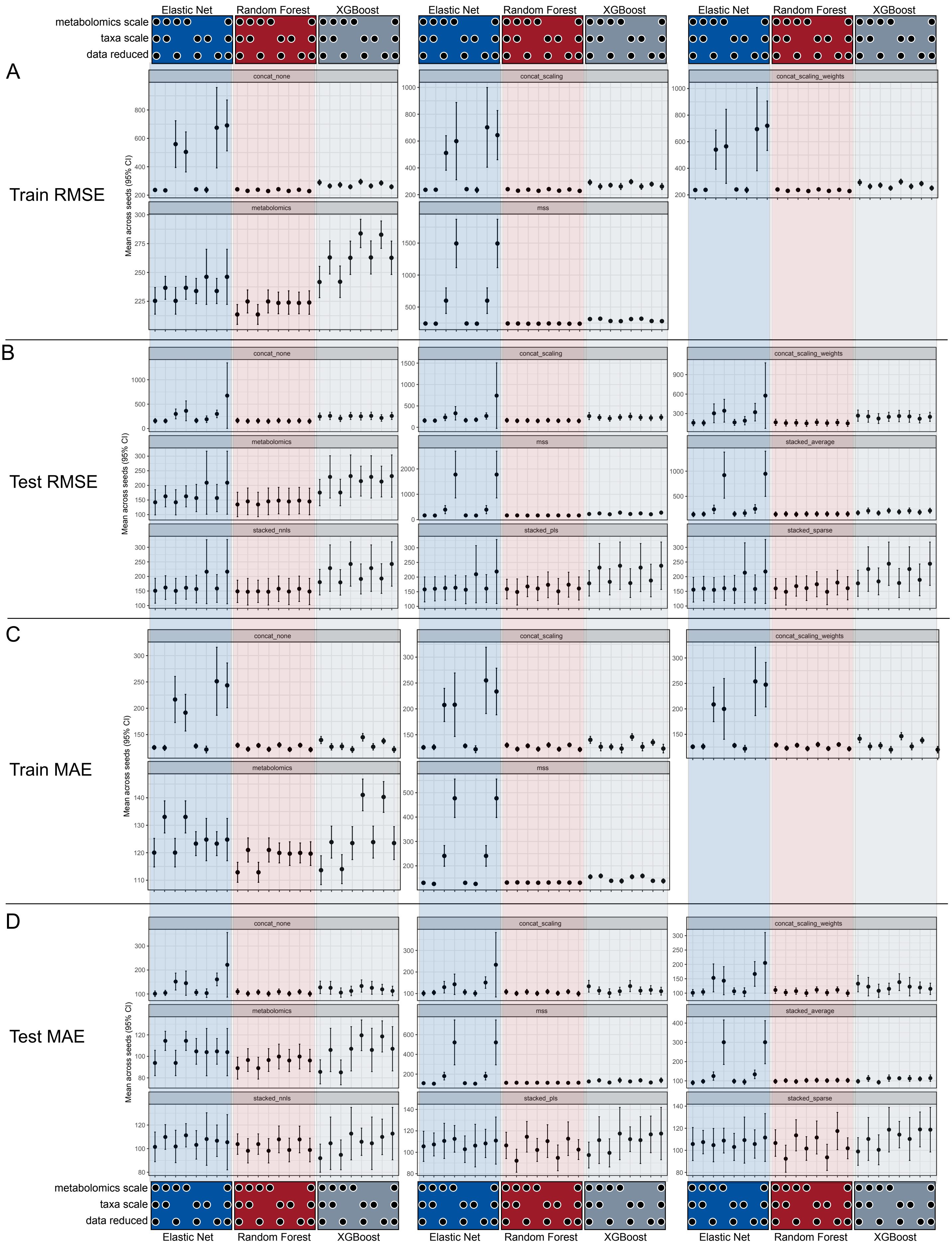

Franzosa Control vs. UC Mean Performance with 95% CI Across Seeds
(● Data is scaled and/or reduced)

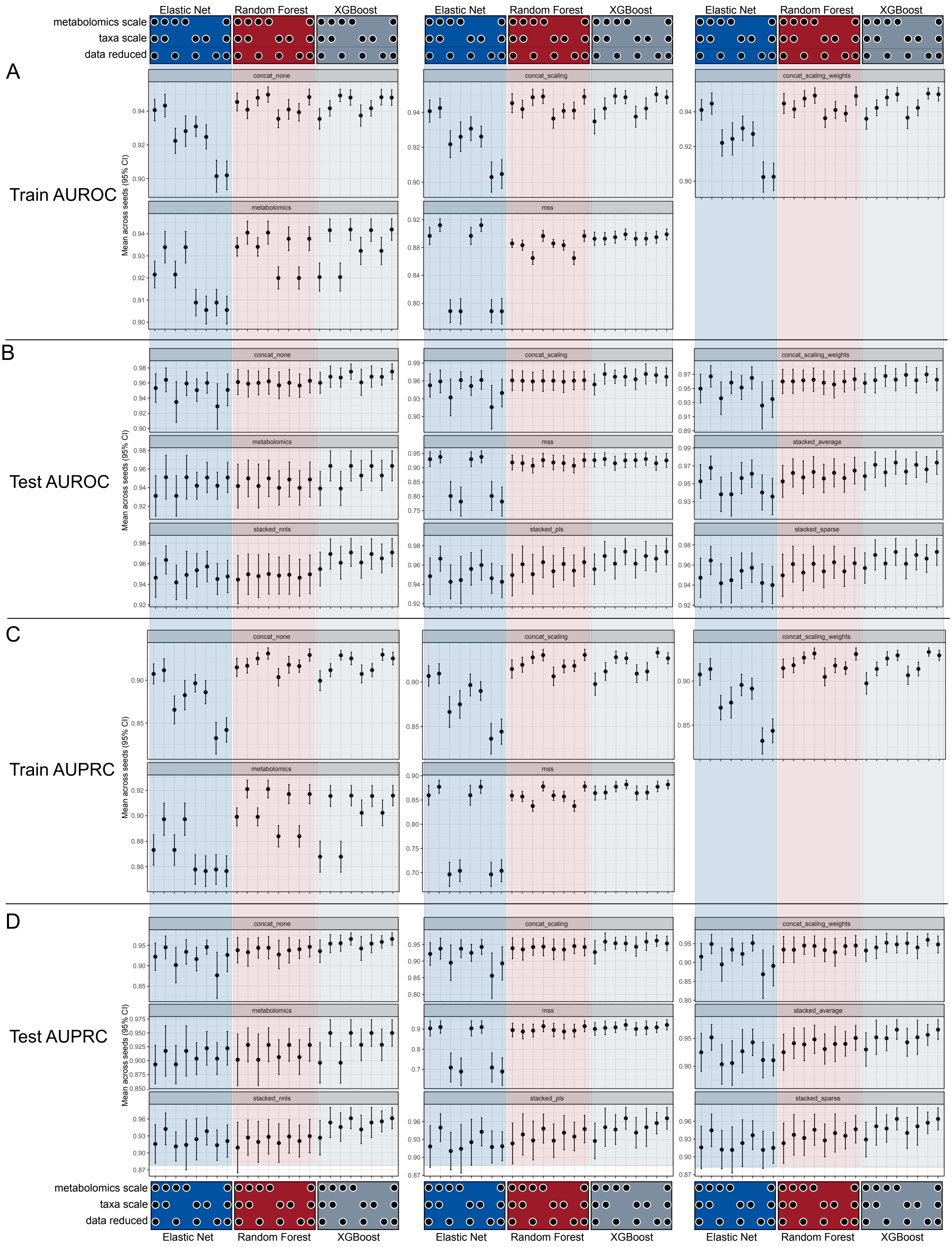

Franzosa UC Fp Mean Performance with 95% CI Across Seeds (● Data is scaled and/or reduced)

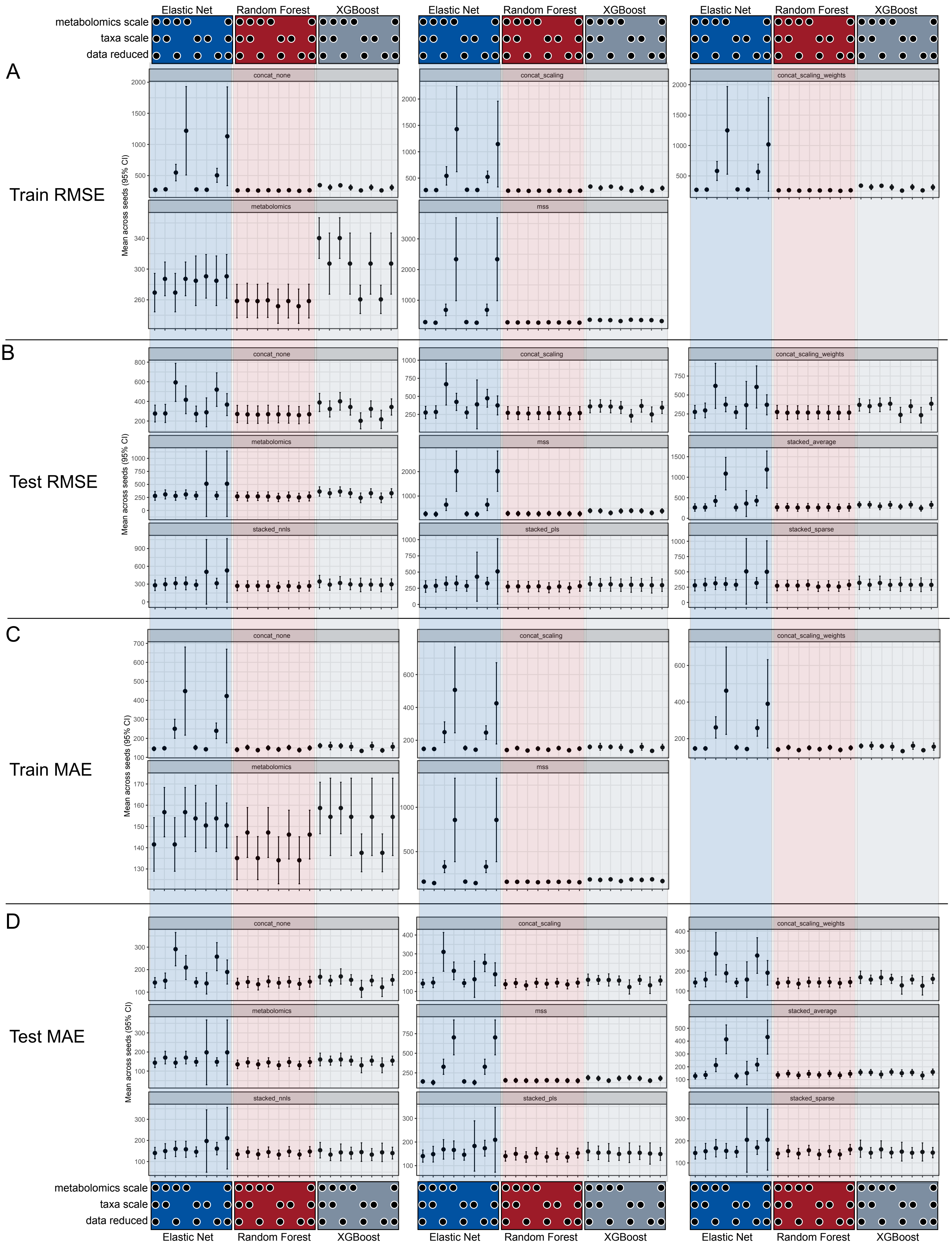

Wang Control vs. Kidney Failure Mean Performance with 95% CI Across Seeds
(●Data is scaled and/or reduced)

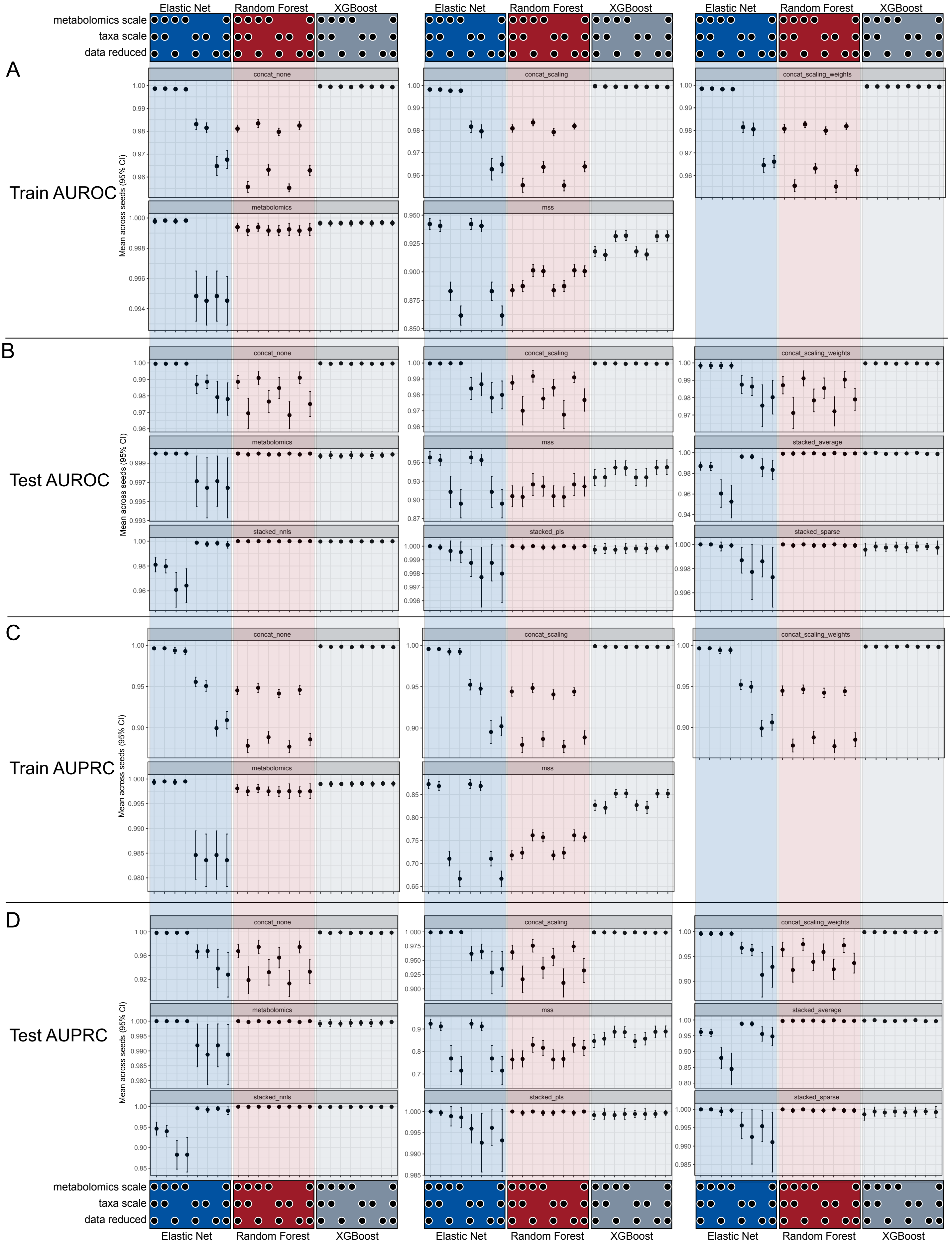

Wang Creatinine Mean Performance with 95% CI Across Seeds (● Data is scaled and/or reduced)

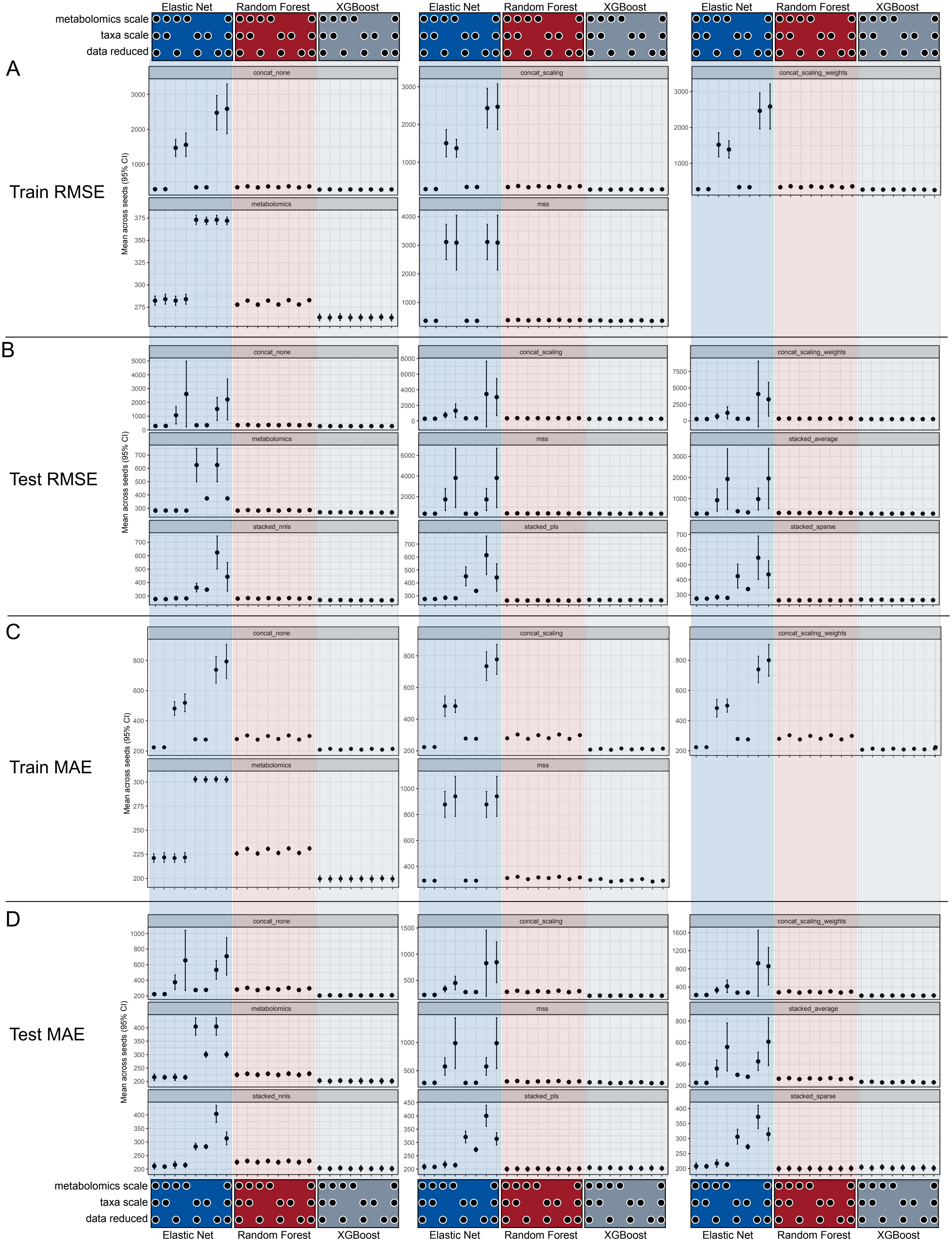

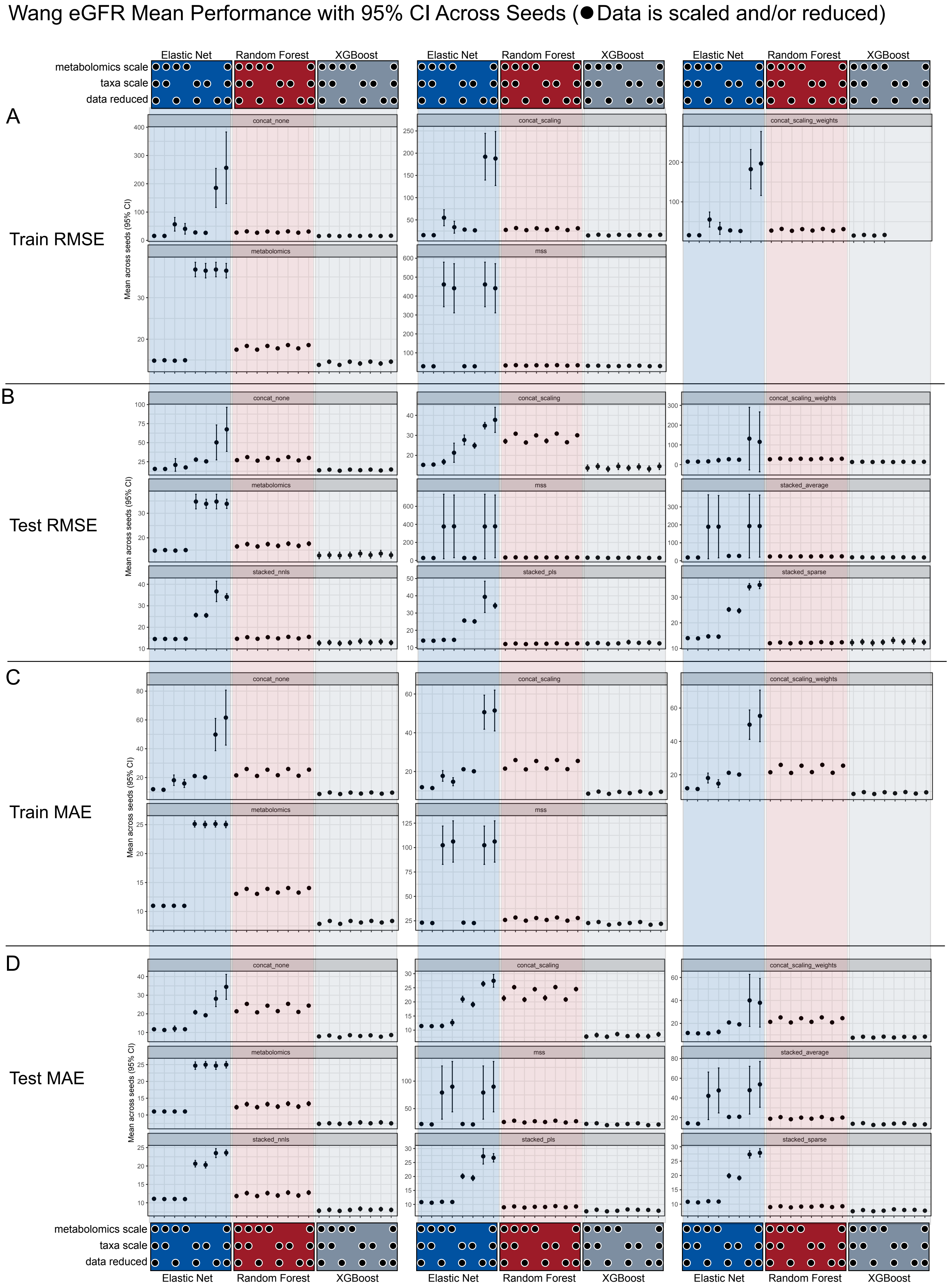

Wang eGFR Mean Performance with 95% CI Across Seeds (● Data is scaled and/or reduced)
A
Train RMSE
B
Test RMSE
C
Train MAE
D
Test MAE
Elastic Net
Random Forest
XGBoost
metabolomics scale
taxa scale
data reduced
concat_none
concat_scaling
concat_scaling_weights
metabolomics
mss
stacked_average
stacked_nnls
stacked_pls
stacked_sparse
Mean across seeds (95% CI)

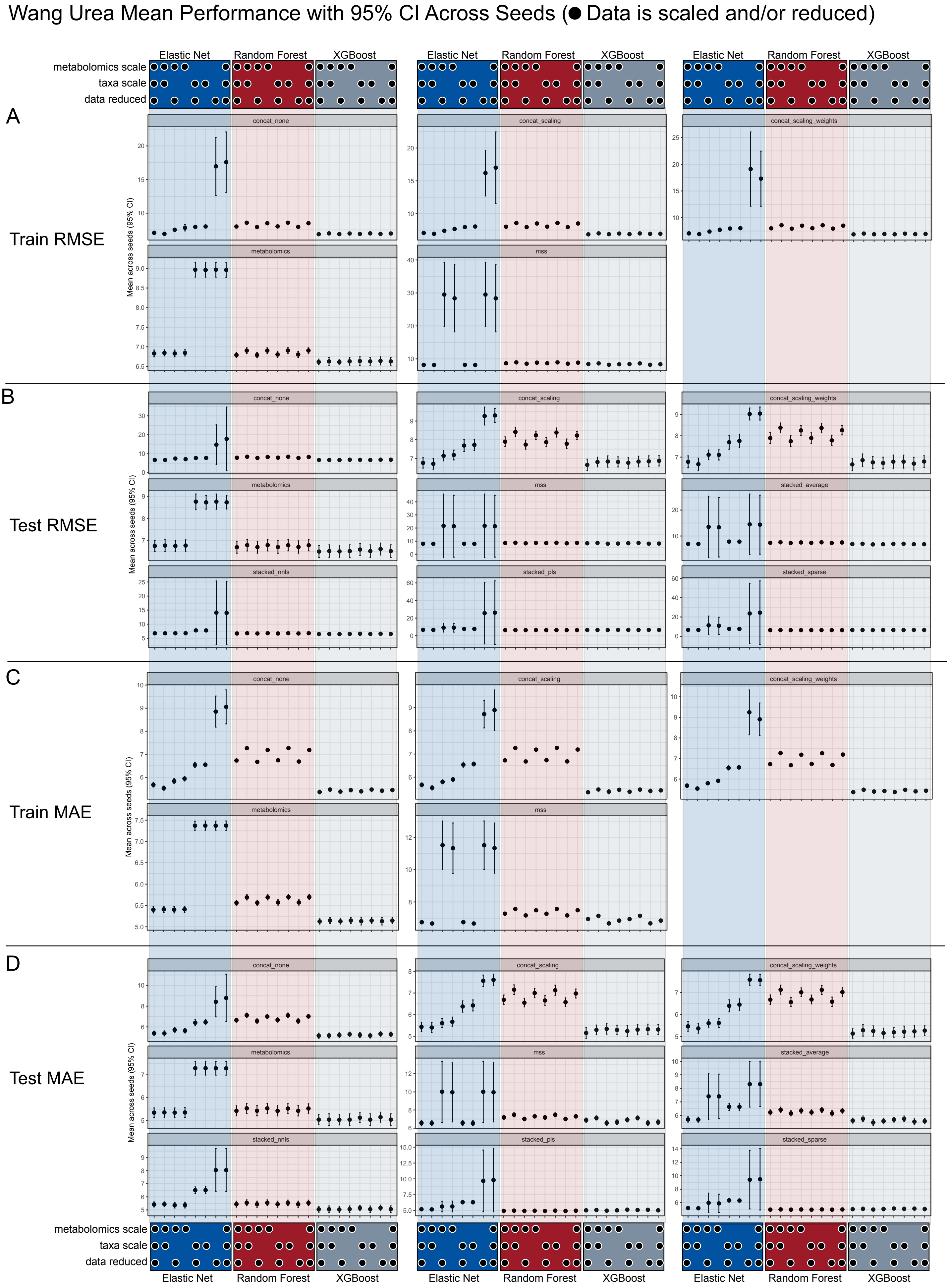

Wang Urea Mean Performance with 95% CI Across Seeds (● Data is scaled and/or reduced)
Elastic Net
Random Forest
XGBoost
metabolomics scale
taxa scale
data reduced
A
Train RMSE
B
Test RMSE
C
Train MAE
D
Test MAE
concat_none
concat_scaling
concat_scaling_weights
metabolomics
mss
stacked_average
stacked_nnls
stacked_pls
stacked_sparse
Mean across seeds (95% CI)

Yachida Control vs. MP & CRC Stage 0 Mean Performance with 95% CI Across Seeds
(● Data is scaled and/or reduced)

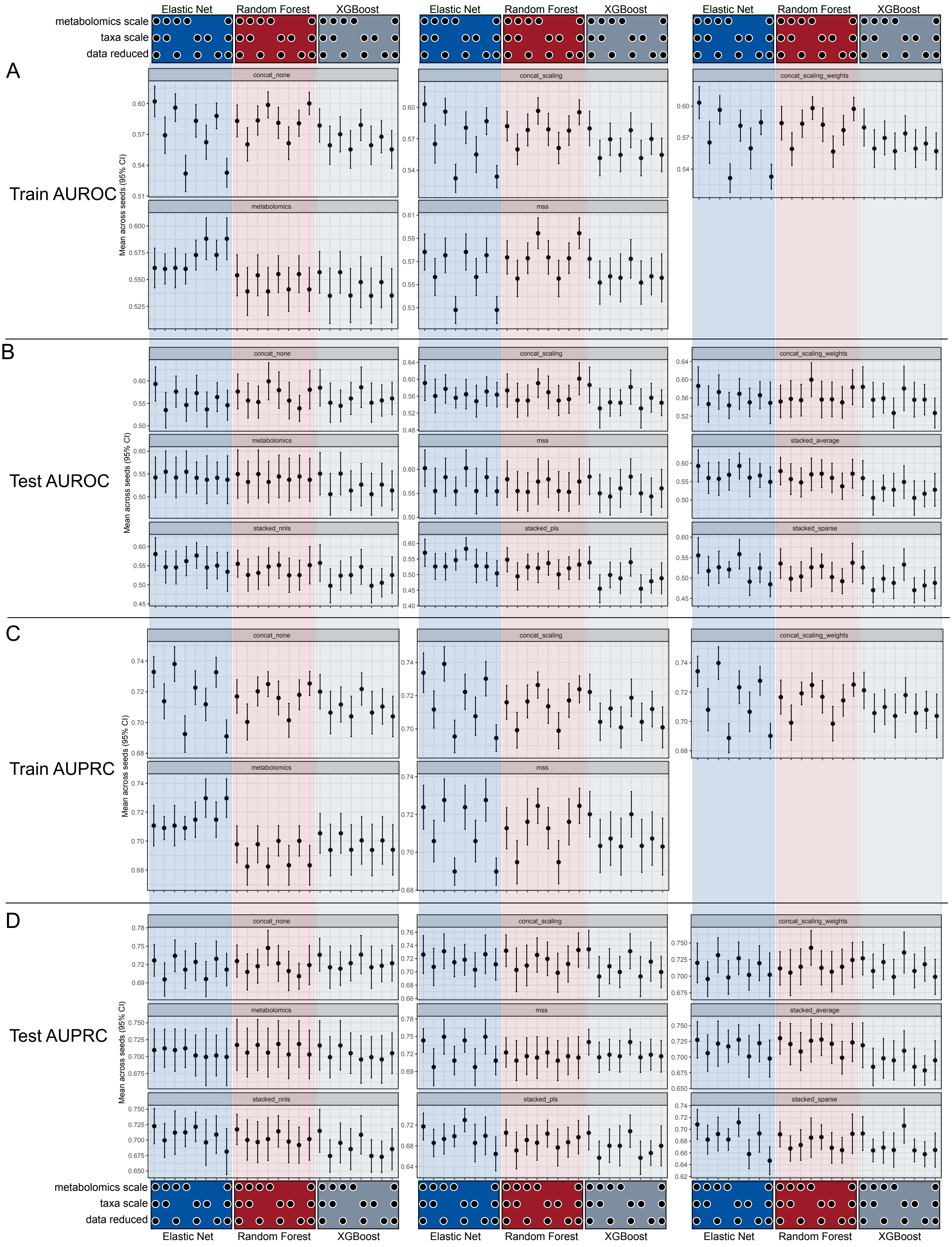

Yachida Control vs. CRC Stages 1 & 2 Mean Performance with 95% CI Across Seeds
(● Data is scaled and/or reduced)

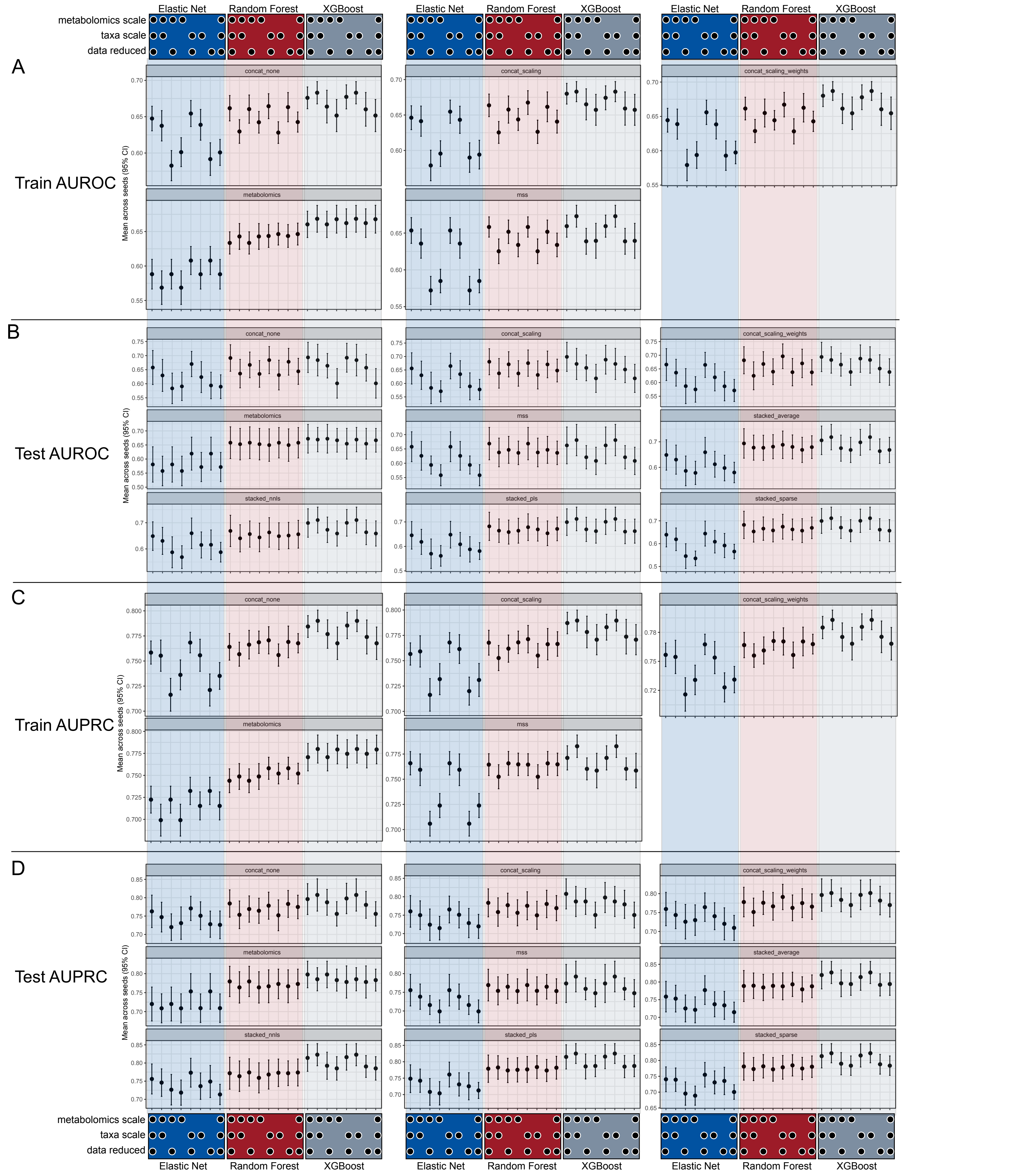

### Yachida Control vs. CRC Stages 3 & 4 Mean Performance with 95% CI Across Seeds
(● Data is scaled and/or reduced)

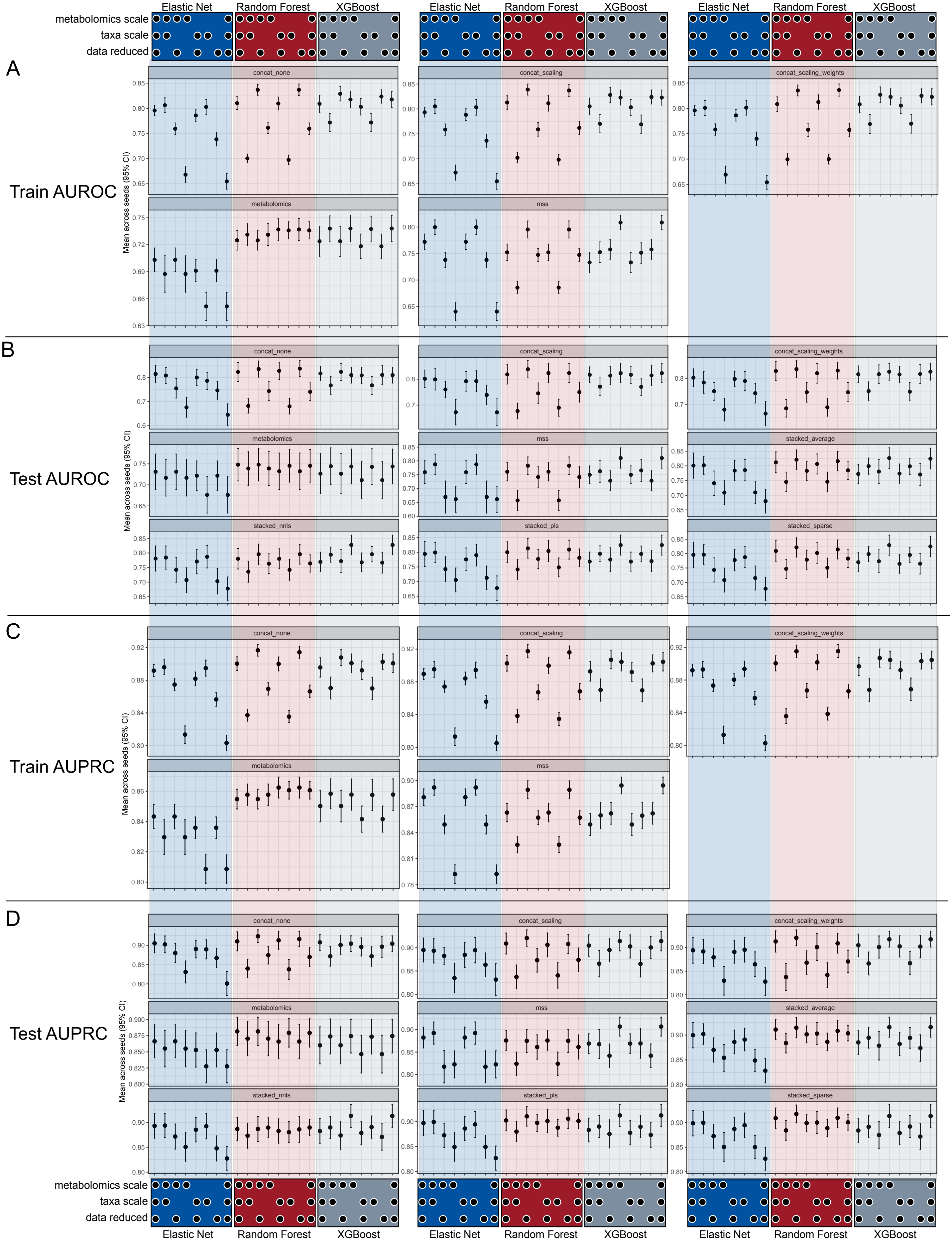

Yachida Control vs. CRC All Stages Mean Performance with 95% CI Across Seeds
(●Data is scaled and/or reduced)

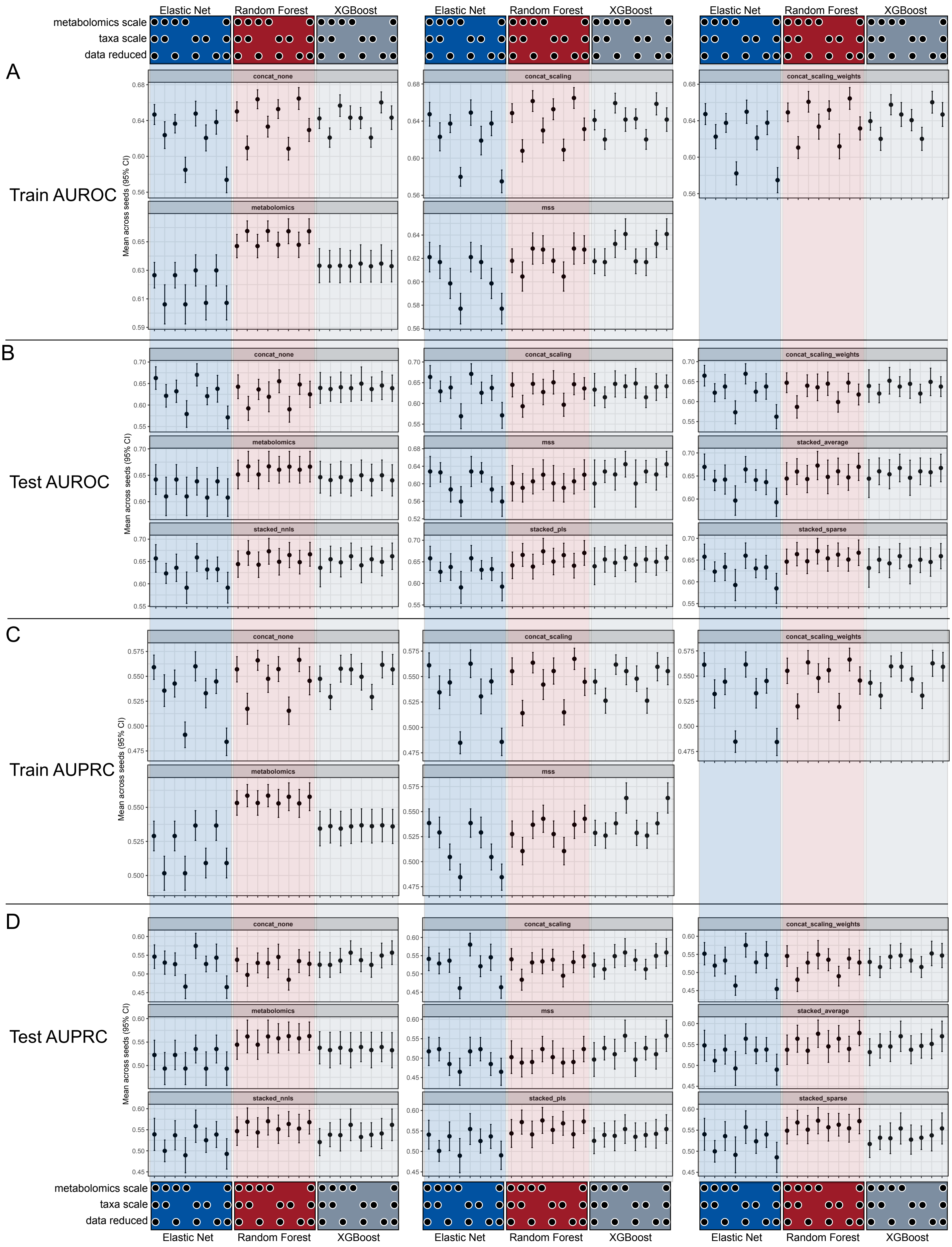
